# *In vivo* engineered B cells retain memory and secrete high titers of anti-HIV antibodies in mice

**DOI:** 10.1101/2021.04.08.438900

**Authors:** A. D. Nahmad, C. R. Lazzarotto, N. Zelikson, T. Kustin, M. Tenuta, D. Huang, I. Reuveni, M. Horovitz-Fried, I. Dotan, R. Rosin-Arbesfeld, D. Nemazee, J.E. Voss, A. Stern, S. Q. Tsai, A. Barzel

## Abstract

As a potential single-shot HIV therapy, transplanted engineered B cells allow robust secretion of broadly neutralizing antibodies (bNAbs). However, *ex vivo* engineering of autologous B cells is expensive and requires specialized facilities, while allogeneic B cell therapy necessitates MHC compatibility. Here, we develop *in vivo* B cell engineering, by injecting two adeno associated viral vectors, one coding for saCas9 and another coding for a bNAb. Following immunizations, we demonstrate memory retention and bNAb secretion at neutralizing titers. We observed minimal CRISPR/Cas9 off-target cleavage, using unbiased CHANGE-Seq analysis, while on-target cleavage at undesired tissues is reduced by expressing saCas9 from a B cell specific promoter. *In vivo* B cell engineering is thus a safe, potent and scalable method for expressing desired antibodies against HIV and beyond.

**One sentence summary:** B cells can be engineered *in vivo* to robustly secrete anti-HIV bNAbs in a safe, durable and scalable manner.

## INTRODUCTION

Broadly neutralizing antibodies (bNAbs) against HIV can suppress the viremia. In particular, combination therapy with the bNAbs 3BNC117 and 10-1074 allowed long-term suppression upon interruption of antiretroviral therapy (ART) in individuals with antibody-sensitive viral reservoirs^1^. Similarly, viremic individuals with dual antibody-sensitive viruses, experienced diminished viremia for three months following the first of up to three dual-bNAb infusions^2^. However, the mean elimination half-life of the bNAbs is 16 and 23 days, respectively^3^, allowing the virus to rebound. Moreover, individuals with prior resistance to one of the bNAbs have mounted resistance to the second antibody, and individuals with prior resistance to both antibodies were excluded from the trials. Limited bNAb persistence may be addressed by constitutive expression from muscle following viral vector transduction^4,5^. However, anti-drug antibodies (ADA) may develop^6^, possibly because of improper glycosylation. Moreover, antibodies expressed from muscle do not undergo class switch recombination (CSR) or affinity maturation, which may be required for long term suppression of a diverse and continuously evolving HIV infection. In order to overcome these challenges, we^7,8^ and others^9–13^ have developed B cell engineering for antibody expression. In particular, we previously combined Toll-like receptor (TLR)-mediated *ex vivo* activation of B cells with *in vivo* prime-boost immunizations, and demonstrated that engineered B cells allow immunological memory, CSR, somatic hypermutation (SHM) and clonal selection. However, cost and complexity of autologous B cell engineering *ex vivo* may be prohibitive. At the same time, use of engineered allogeneic B cells is challenging due to the requirement for HLA matching for receiving T cell help and avoiding graft rejection. As an alternative, we describe here an *in vivo* B cell engineering protocol, a safe and scalable method based on a single systemic injection of adeno associated viral (AAV) vectors coding for CRISPR/Cas9 and for the desired bNAb cassette, which is targeted for integration at the IgH locus.

## RESULTS

In order to promote *in vivo* B cell engineering, we used a pair of AAV-DJ vectors^14^, one coding for saCas9^15^ and the other coding for the 3BNC117 anti-HIV bNAb^16^ (Fig. 1A). In the first set of experiments, the saCas9 is expressed from the ubiquitously active CMV promoter, and the sgRNA, targeting saCas9 to the IgH locus, is coded on the same AAV. The bNAb, in turn, is coded as a bi-cistronic cassette under the control of an IgH-enhancer-dependent promoter and flanked by homology arms to the desired saCas9 cut-site within the J-C intron of the IgH locus (Fig. S1^7^). The bNAb cassette includes the full light chain and the variable segment of the heavy chain (V_H_) being separated by a sequence coding for a Furin cleavage site and for a 2A-peptide. A splice donor sequence follows the V_H_ gene segment in order to allow its fusion to constant IgH exons, upon integration into the locus and subsequent transcription and splicing. Our design facilitates disruption of the endogenous IgH locus and initial bNAb expression as a membranal B cell receptor (BCR). Importantly, this allows for subsequent activation of the engineered B cells upon antigen-binding, which leads to differentiation into memory and plasma cells.

**Fig. 1:**
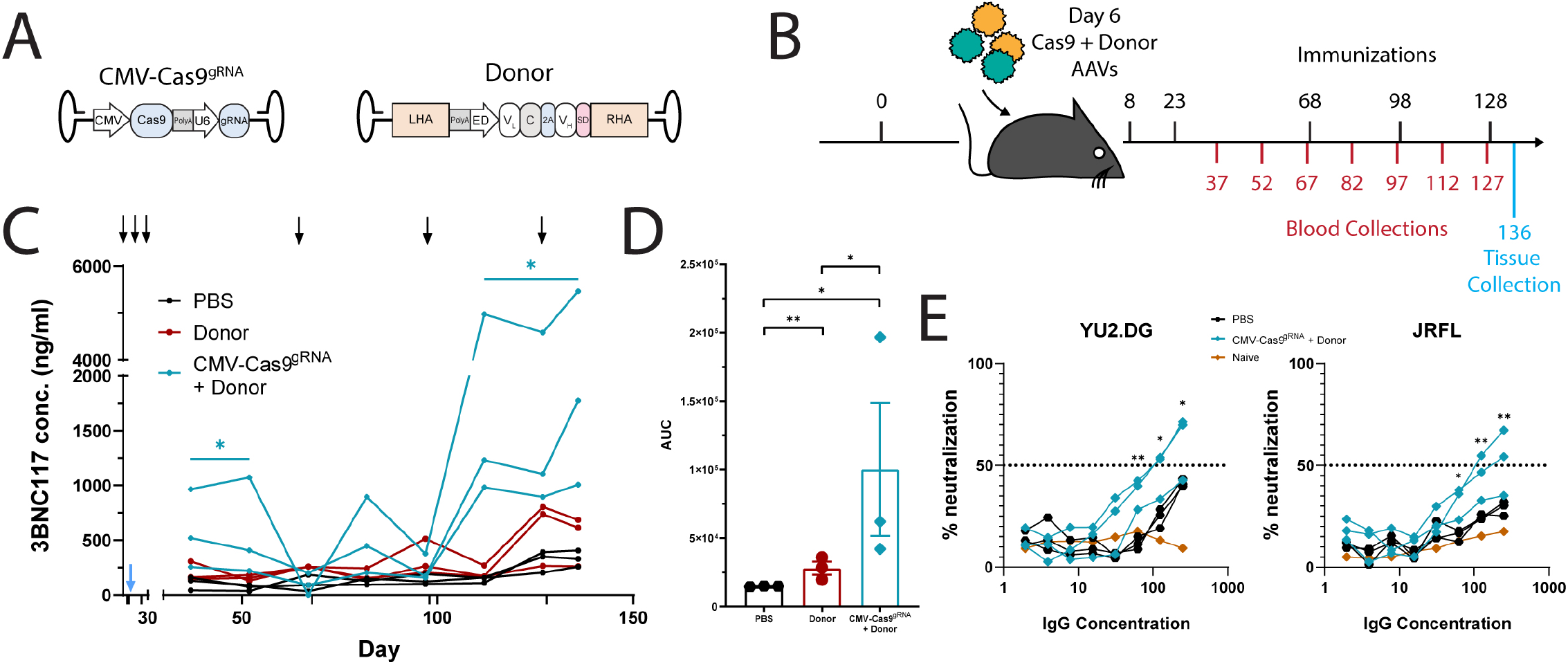
*In vivo* engineering of B cells to express an anti-HIV bNAb. **A**. Vector design. The expression of saCas9 and the sgRNA is driven by the CMV and U6 promoters, respectively^15^. The Donor vector encodes the light chain (V_L_-C), and the variable heavy chain (V_H_) of the 3BNC117 antibody under the regulation of an enhancer dependent (ED) promoter. A splice donor (SD) sequence downstream to the V_H_ segment facilitates splicing with the endogenous constant segment upon integration into the IgH locus (Fig. S1), while an upstream polyadenylation (polyA) signal prevents undesired splicing between the endogenous V_H_ and constant segments^7^. The donor cassette is targeted by homology directed repair (HDR) to the IgH locus, using the flanking homology arms (LHA and RHA) **B**. Experimental scheme. Immunizations are indicated in black, above the timeline. Blood collections are indicated in red, below the timeline. A single injection of AAVs occurs at day 6, with 5E11 vg per mouse for each vector. **C**. 3BNC117 IgG titers as quantified by ELISA using an anti-idiotypic antibody to 3BNC117. The black arrows indicate immunizations and the blue arrow indicates the AAV injection. Each line represents a mouse. * = pv<0.05, two-way ANOVA of CMV-Cas9^gRNA^ + Donor as compared to the Donor group. n=3. **D**. Area under the curve (AUC) of C. * = pv<0.05, ** = pv<0.01 unpaired t-test. n=3. **E**. Transduction neutralization of TZM.bl cells by the YU2.DG (left) and JRFL (right) HIV pseudoviruses in the presence of IgGs purified form day 136 sera. Neutralization is calculated as percent reduction from maximal luminescence per sample. The PBS control received immunizations as in C, while the naïve control represents serum IgG from an untreated mouse. * = pv<0.05, ** = pv<0.01 Two-way ANOVA with Šidák’s multiple comparison for time points comparison to PBS. AUC bar graphs are available in Fig. S5.

B cell activation is required for efficient AAV transduction^17^. Subsequent activation signals for the engineered B cells may benefit from prior priming of T helper cells and from presentation of appropriate immune complexes by follicular dendritic cells^18^. AAV injections to mice were thus preceded by pre-immunizations, modeling a pre-existing infection. In particular, C57BL/6 mice were immunized with 20 µg of the gp120 HIV antigen, which is the target of 3BNC117. On day 6 post-immunization, each mouse was injected with 5E11 vg of a bNAb coding (donor) vector, 5E11 vg of the saCas9 coding vectors, or both (Fig. 1B). The mice then received additional immunizations at days 8, 23, 68, 98 and 128. Following the boosting regimen, mice receiving both a donor vector and an saCas9 vector had up to 5 µg/ml of the 3BNC117 bNAb in their blood (Fig. 1C-D). While this experiment used the monomeric gp120 antigen of the clade B HIV strain YU2.DG, high titers could also be obtained in an independent experiment using the clade A, BG505-based native trimer nanoparticle immunogen (MD39-ferritin)^19^, attesting for the breadth of the 3BNC117 expressing cells *in vivo* (Fig. S2). Mice injected with both a donor vector and an saCas9 vector had much higher 3BNC117 titers than mice receiving donor vector only. Nevertheless, 3BNC117 titers in mice receiving only the donor vector exceeded the background levels measured in mice injected with PBS (Fig. 1D). Indeed, integration of the antibody gene into the IgH locus was evident by nested RT-PCR followed by Sanger sequencing on splenic B cell RNA from mice receiving dual vector injection as well as from two of the mice injected with the donor vector only (Fig. S3). Notably, engineered cells have secreted antibodies of multiple isotypes (Fig. S4). Finally, IgG purified from treated mice can neutralize autologous YU2.DG and heterologous JRFL HIV pseudoviruses (Fig. 1E, S5). This demonstrates a potent and functional bNAb response from *in-vivo* engineered B cells upon vaccination.

Flow cytometry was used to estimate the engineering rates at the cellular level. The frequency of 3BNC117-expressing cells reached 1-3% of total blood B cells following the later immunizations in all mice injected with both a bNAb vector and an saCas9 vector, but not in mice injected with PBS or with the bNAb vector alone (Fig. 2A-C). Upon sacrificing the mice at day 136, 8 days after the last immunization, up to 23% of the plasmablasts in the spleen expressed 3BNC117 (Fig. S6). In addition, the rate of 3BNC117-expressing splenic cells was between 5% to 10% among germinal center (GC) B cells (Fig. 2D-E, S7), while between 1% to 2% of bone marrow lymphocytes were 3BNC117-expressing B cells (Fig. S8). No significant expression of 3BNC117 was detected in mice receiving donor vector alone in any of the tissues. In order to study somatic hypermutation and clonal selection, we extracted DNA from the liver and the spleen of one of the treated mice at day 136 and performed Illumina sequencing of amplified 3BNC117 V_H_ segments. Much of the mutation repertoire was shared between the liver and the spleen and may thus reflect heterogeneity in AAV production that is subjected to little or no selection. In particular, all the 3BNC117 V_H_ variants found to be over-represented in the liver are also over-represented in the spleen. Importanlty however, the inverse is not true. The CDR1 substitution R30K is the most prevalent substitution in the spleen. It accounts for more than 20% of all mutants in the spleen but is found at very low abundance in the liver (Fig. 2F). Indeed, bNAbs of the VRC01 family were shown to have side chain interactions with the HIV gp120 antigen at position 30^16^ One may speculate that the conservative R30K substitution in 3BNC117 relieves some steric clash upon binding to monomeric gp120. Including R30K, a total of five different positions along the V_H_ segment showed signs of positive selection in the spleen by dn/ds analysis (Fig. S9), although analysis may be less informative regarding the two positions at the distal extremes, which are assigned high dn/ds values also in the liver. We conclude that our *in vivo* engineering and immunization scheme has led to clonal expansion that is limited in span but pronounced in its magnitude.

**Fig. 2:**
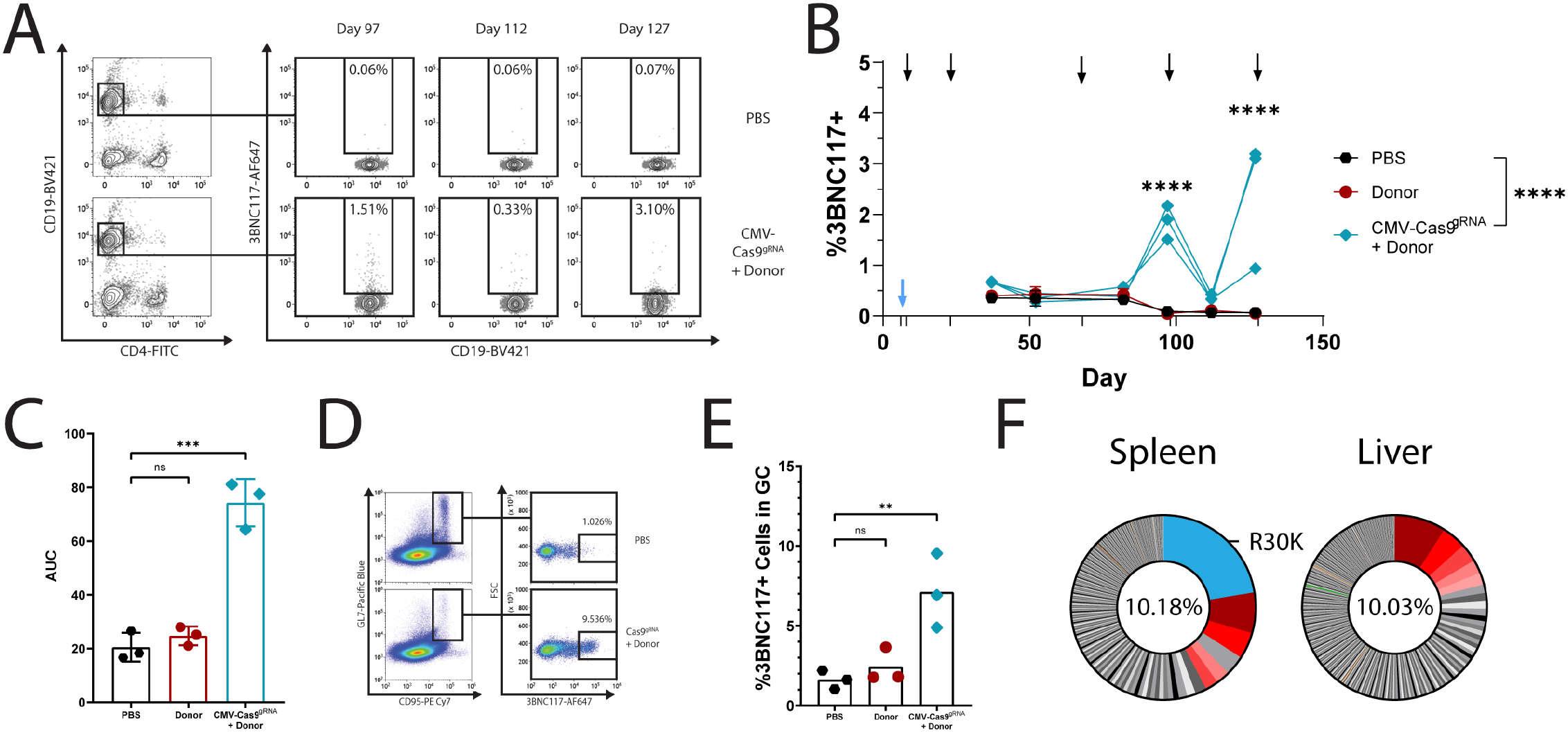
*In-vivo* engineered B-cells are found in lymphatic tissues 130 days following AAV injection. **A**. Flow cytometry plots demonstrating 3BNC117 expression among blood B cells (CD19^+^, CD4^-^). **B**. Quantification of blood 3BNC117-expressing cells over time. The black arrows indicate immunizations and the blue arrow indicates AAV injection. **** = pv<0.0001 Two-way ANOVA with Šidák’s multiple comparison for time points comparison to PBS. Each line represents a mouse.**C**. Area under the Curve (AUC) analysis of the data presented in B. **D**. Flow cytometry plots demonstrating 3BNC117 expression of cells with a germinal center phenotype (GL7^+^, CD95/Fas^+^) in the spleen. Pre-gated on live, singlets. **E**. quantification of D. Mean is indicated by the bars, ns= non-significant, ** = pv<0.01 One way ANOVA with Tukey’s multiple comparison. **F**. Pie charts of 3BNC117 V_H_ variants amplified from spleen and liver DNA at day 136. Red shading indicates expanded variants that are shared between the liver and the spleen. Blue shading indicates the R30K variant. Numbers in the middle of the pies indicate the total frequency of mutant reads in these samples.

Next, we assessed the possible off-target effects of our *in vivo* engineering approach. First, we quantified the copy number of the bNAb cassette from various tissues. Then bNAb cassette was found at a high copy number in the liver at day 37 (Fig. 3A) and the levels were reduced by only 10 fold at day 136 (Fig. 3B), reflecting high retention of AAV episomes in the liver. High copy number was also found in the blood at day 37, but levels dropped sharply by day 136, perhaps due to multiple cell divisions (Fig. S10A). Interestingly, the AAV copy number in the bone marrow was significantly increased from day 37 to day 136, and a non-significant similiar trend was also detected in the lymph nodes, indicating the possible accumulation of 3BNC117-expressing cells in these tissues (Fig. 3B). The copy number in the liver was similar whether or not the saCas9 coding AAV was co-injected to the mice. In contrast, donor AAV copy number in the lymph nodes and in the bone marrow was found to be logs higher with saCas9 AAV co-injection, signifying the selection of 3BNC117-expressing B cells (Fig. 3C, S10B). To define the genome-wide off-target activity of SaCas9, we performed circularization for high-throughput analysis of nuclease genome-wide effects by sequencing (CHANGE-seq)^20^ on genomic DNA from C57BL/6 mice. 31 on- and off-target sites were identified, characterizing the sgRNA target site as highly specific (CHANGE-seq specificity ratio = 0.95) (Fig. 3D). We then performed targeted sequencing on four off-target sites and the on-target site using genomic DNA from liver and spleen of treated mice and from a negative control mouse. Relative to control DNA from the spleen of an untreated mouse, a trend for a higher mutation rate, indicating error prone repair of CRISPR/Cas9 induced double-stranded DNA breaks, was evident at the IgH on-target site but not in any of the tested off-target sites (Fig. 3E).

**Fig. 3:**
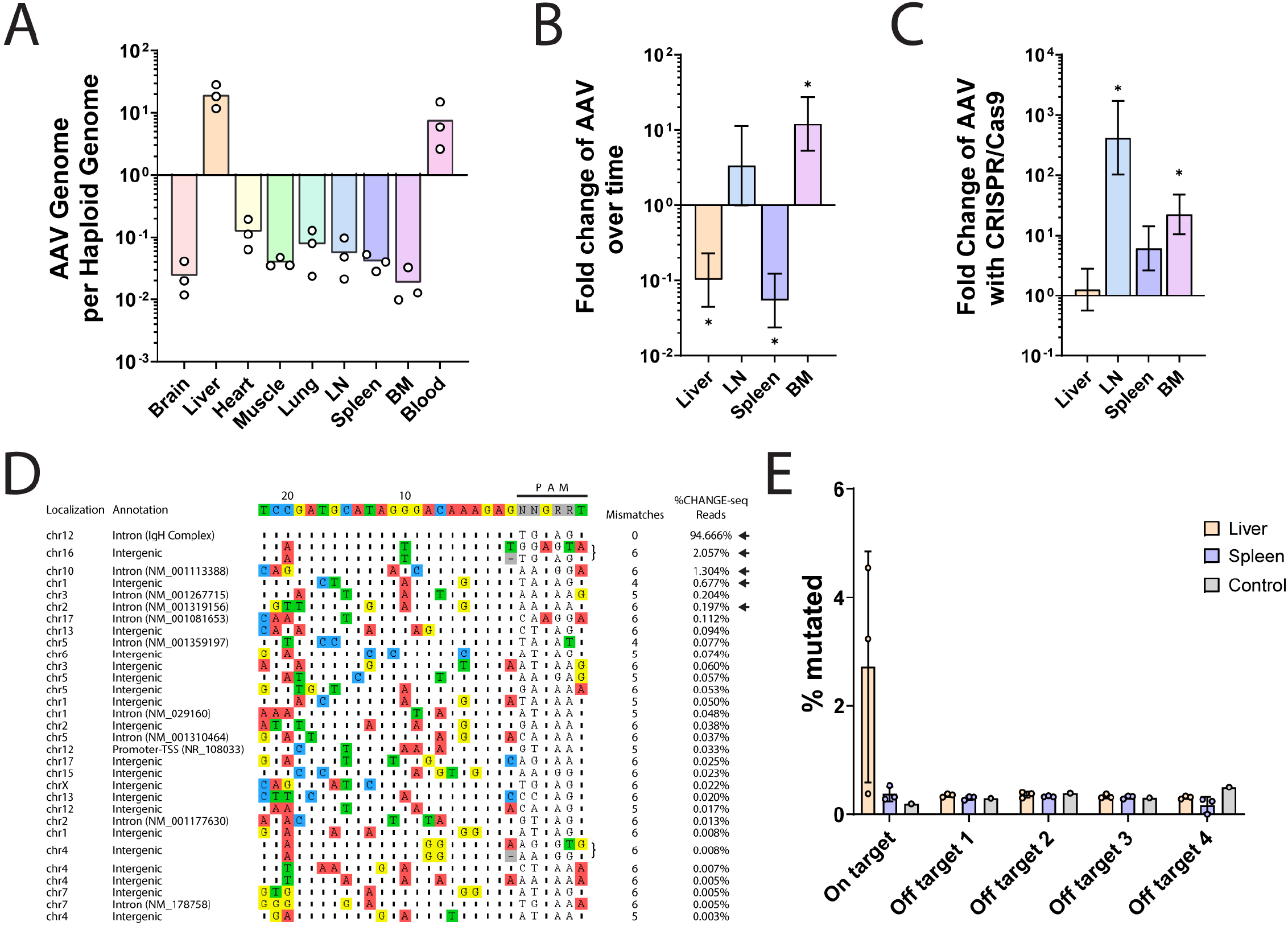
AAV biodistribution and saCas9 off-target cleavage analysis reveal a high safety profile. **A**. Donor AAV copy number quantification by qPCR in indicated tissues at day 136 in mice injected with two AAVs as in Fig. 1A. **B**. Relative copy number of donor AAV between day 37 and day 136 in selected tissues. **C**. Relative copy number of donor AAV between mice injected with two AAVs as in figure 1A and mice injected with donor AAV only, at day 136. For B. and C. Mean, upper and lower boundaries are indicated. * = pv<0.05 for comparison between the two time points or mice groups, respectively, unpaired t-test. n=3.Y axis in A-C uses a log scale. **D**. Unbiased CHANGE-seq analysis of potential saCas9 off-target cleavage with the sgRNA used in this study. Localization, annotation in the genome, number of mismatches and % read counts are indicated for each on- or off-target site. Sequence of the sgRNA with the PAM is indicated on the top. Black arrows indicate target sites used for *in-vivo* analysis. Mismatches between off-target sites and intended sgRNA target are color-coded. **E**. On- and off-target saCas9 cleavage of target sites indicated in D. by black arrows, in the spleen (mauve) and liver (beige) of mice injected with two AAVs as in figure 1A at day 136 as compared to uncut splenic lymphocytes DNA. Each dot represents a mouse.

The coding of the sgRNA together with the saCas9 on the same AAV is predicted to allow DNA cleavage in many cells that are not co-transduced with the donor AAV. The resulting, non-productive, cleavage may be avoided if the sgRNA cassette is instead separated from saCas9 gene and coded on the donor AAV (Fig. S11). Repeating the above mouse experiments (Fig. 1B) with this new pair of AAVs allowed high 3BNC117 titers following repeated immunizations (Fig. S12), and the frequency of 3BNC117-expressing cells reached 0.5%-3% of total blood B cells (Fig. S13). Upon sacrificing the mice at day 136, up to 10% of splenic plasmablasts expressed 3BNC117 (Fig. S14). In addition, up to 7% of splenic B cells with a germinal center phenotype expressed 3BNC117 (Fig. S15), while 3% of the bone marrow lymphocytes were 3BNC117-expressing B cells (Fig. S16). These results are of the same range as those obtained when the sgRNA was coded together with the saCas9, although direct side by side comparison is hindered by the use of different ubiquitously active promoters. Importantly, the overall numbers of splenic plasmablasts, germinal center B cells and bone marrow plasma cells were similar to those in the control groups (Fig. S17), mitigating concerns of B cell neoplasm. In order to further increase the safety of our approach, we next coded the saCas9 under the control of the CD19, B cell specific, promoter^21^ (Fig. 4A). In particular, C57BL/6 mice were immunized with 20 µg of HIV gp120, and 6 days later each mouse was co-injected with one vector coding for the bNAb and for the sgRNA and with a second vector coding for saCas9, regulated by the CD19 promoter (Fig. 4A-B). The mice then received additional immunizations at days 8, 23, 38, and 53. Following the boosting regimen, treated mice had up to 2 μg/ml of the 3BNC117 bNAb in their blood (Fig. 4C), attesting that replacing the promoter did not preclude the therapeutic effect. Different groups of mice were sacrificed for on-target cleavage analysis 3 days after having been co-injected with the donor + sgRNA vector and with a second vector coding for saCas9 under the control of either the ubiquitously active SFFV promoter or the B cell specific CD19 promoter. Similar transduction rates were obtained for vectors coding the saCas9 under the regulation of the CD19 or SSFV promoters (Fig. S18). However, the CD19 promoter significantly reduced saCas9 expression in the liver, but not in peripheral blood mononuclear cell (PBMCs, Fig. 4D). Importantly, the rates of on-target cleavage in the liver, as measured by TIDE analysis, were significantly above background only when using the SFFV promoter, rather than the CD19 promoter, to drive saCas9 expression (Fig. 4E). Therefore, separating the coding of saCas9 and the sgRNA between the two AAVs and expressing saCas9 under a B cell specific promoter reduce undesired cleavage to below our limit of detection while allowing high 3BNC117 titers following immunizations.

**Fig. 4:**
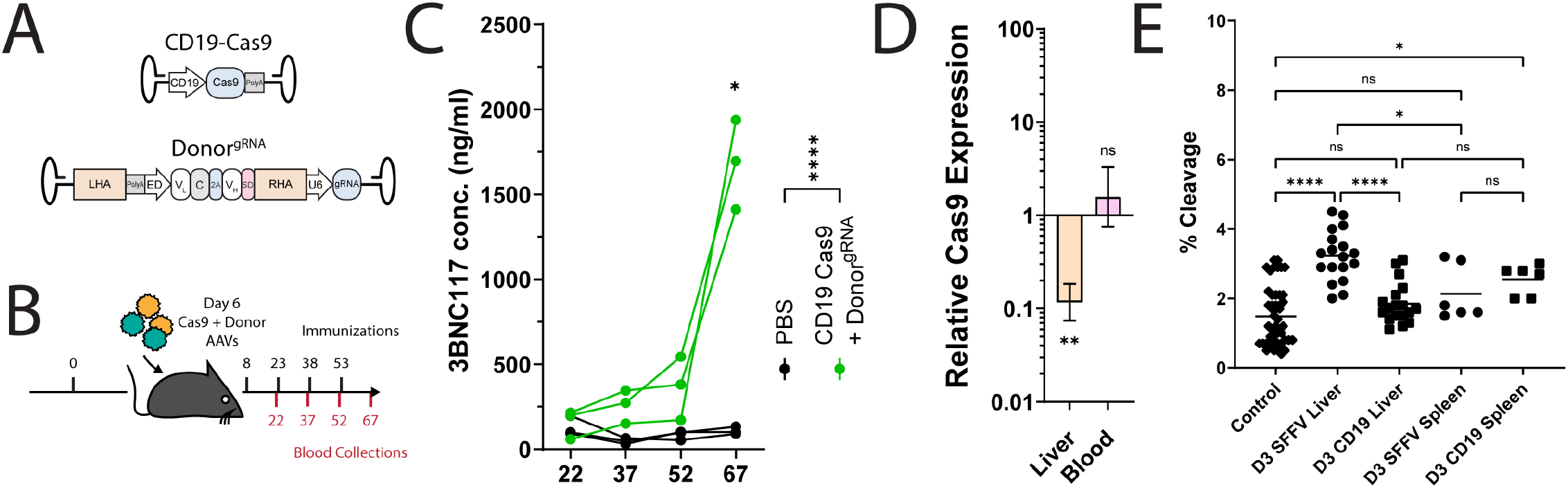
Safety may be improved by coding saCas9 and the sgRNA on separate AAVs and by expressing saCas9 under the regulation of a B-cell specific promoter. **A**. Map of the AAV vectors used. saCas9 is expressed under the CD19 promoter, while the sgRNA is coded on the donor vector, outside of the homology arms. **B**. Experimental scheme. Mice were immunized according to the timeline in black (top), and bled as indicated in red (bottom). **C**. 3BNC117 IgG titers as quantified by ELISA with an anti-idiotypic antibody. Titers peak following the fourth immunization after AAV injection. Each line represents a mouse. * = pv<0.05 and **** = pv<0.0001 for Two-way ANOVA with Šidák’s multiple comparison for time point comparison. **D**. Relative saCas9 expression, depicted as the ratio between saCas9 expression from the CD19 promoter and saCas9 expression from the SFFV promoter. Error bars correspond to lower and upper boundaries derived from unpaired t test. For A. and B., ns = non-significant, ** = pv<0.01. n=3. **E**. TIDE analysis of on-target cleavage in the indicated tissues using either SFFV or CD19 driven saCas9 expression. Control samples come from naive splenic lymphocytes. ns = non-significant, * = pv<0.05 and **** pv<0.0001 One-way ANOVA with Tukey’s multiple comparison. Each dot represents a comparison between a control sample and an independent mouse sample.

Eliciting a specific, neutralizing antibody response to hypervariable viruses is a long-standing challenge in medicine. B cell engineering provides an opportunity to express desired therapeutic antibodies for adaptive immunity. Here, we uniquely demonstrate that B cells can be safely and robustly engineered *in vivo*. A single, systemic dose of dual AAV-DJ coding for CRISPR/Cas9 and donor cassette in mice allowed for site-specific integration, with limited off-target Cas9 expression and DNA double-strand breaks. Upon immunizations, the engineered B cells undergo antigen-induced activation leading to memory retention, clonal selection and differentiation into plasma cells that secrete the bNAb at neutralizing levels. Future modifications may include coding the bNAb as a single chain^13^ to reduce mispairing of the bNAb heavy chain with the endogenous light chain. Such single chain coding can further allow the expression of bi-specific bNAbs, which may be required to provide long-term protection from HIV resurgence^1^. Safety may be further improved by using more specific nucleases^22,23^ and by having the bNAb gene preceded by a splice acceptor rather than by a promoter, to reduce expression from off-target integration^7,12^. Both safety and efficacy may benefit from embedding B cell specific targeting moieties in the AAV vector^24^ or in a non-viral alternative^25^. The therapeutic impact of our approach may best be evaluated in nonhuman primates with HIV-like infections. Finally, *in vivo* B cell engineering may have diverse future applications as it may be used to address other persistent infections as well as to treat autoimmune disease, genetic disorders, and cancer.

## MATERIALS AND METHODS

### Plasmid cloning

For the CMV-Cas9^gRNA^ vector, pX601^15^ (Addgene) was cleaved with BsaI and pre-annealed, phosphorylated (PNK, NEB), sgRNA coding oligo-deoxynucleotides were ligated using T4 DNA Ligase (NEB). For the CD19-Cas9 vector, pAB270^26^ was cleaved using NotI and SpeI (NEB) and an saCas9 coding fragment, amplified from pX601, as well as the murine CD19 promoter, amplified from wild type C57BL/6OlaHsd genomic DNA, were assembled using Hi-Fi DNA Assembly Mix (NEB). For the SFFV-Cas9 vector, pAB270 was cleaved with NotI and SpeI (NEB). The fragment coding the SFFV promoter was amplified from GW175 (Kay Lab, Stanford) and the saCas9 was amplified from pX601. The fragments were assembled using Hi-Fi DNA Assembly Mix (NEB). For the Donor^gRNA^ vector, the U6-gRNA fragment was amplified from ligated pX601 with the murine IgH sgRNA used in this study, and the fragment was assembled using Hi-Fi DNA Assembly Mix (NEB) into the donor vector pADN171XS^7^, following cleavage with SpeI (NEB). A list of primers used for cloning can be found in Supplementary Table 1. All fragments for cloning were amplified using PrimeStar MAX (Takara).

### rAAV production and injections

rAAV-DJ were produced in HEK293T cells (ATCC) by triple transient transfection using polyethylenimine (PEI, Polysciences Inc). For each vector, fourteen 15 cm dishes were transfected at 80% confluency with pAd5 (helper plasmid), rAAV-DJ genome plasmid and vector plasmid at a 3:1:1 ratio^27^. In total, each plate was transfected with 41.25 μg of DNA. Purification was performed with AAVpro Extraction Kit (Takara) according to the manufacturer protocol. Titer quantification was performed by qPCR using SYBRGreen (PCR Biosystems). A list of primers used for AAV titer quantification can be found in Supplementary Table 1.

### Mouse studies

6-10 weeks old C57BL/6OlaHsd (Envigo) mice were housed and kept at ambient temperature of 19-23°C, humidity of 45-65% and with a 12 h light/12 h dark cycle. Immunizations with gp120-YU2 or MD39-ferritin were performed as previously described, using 20 μg/mouse of antigen in Alum (Invitrogen)^7,8^. For AAV injections, mice were anesthetized with 0.1 mg/g and 0.001 mg/g Ketamine and Xylazine, respectively, and were injected by 5E11 vg/vector/100μl/mouse in PBS. Blood samples from mice were collected in heparin. Cells and serum were separated by centrifugation. Serum was collected from the supernatant. For spleens, whole spleens were extracted from mice and mechanically crushed in PBS to be filtered in a 70 μm cell strainer (Corning). For bone marrow, cells were flushed from the posterior femur and tibia. For blood, spleen and bone marrow, cells were processed with red blood cell lysis buffer (Biolegend) and plated in 1640 RPMI (Biological Industries) supplemented with 10% HI FBS (Biological Industries) until processing. Muscle tissue was processed from femoral muscles. Right or left lung were processed for pulmonic tissue. Right or left hemispheres were processed for brain tissue. Lobes were processed for liver tissue and whole heart was used for cardiac tissue. For lymph nodes, inguinal and cervical lymph nodes were pooled for processing.

### Illumina sequencing and analysis

Total genomic DNA was extracted from tissues using Gentra PureGene Tissue Kit (Qiagen). Initial PCR amplification and the subsequent barcoding PCR reaction of the 3BNC117 V_H_ fragments or the Off Target sites was performed using the proofreading PrimeStarMAX Polymerase (Takara) for 35 cycles and 8 cycles, respectively. A list of primers used for these reactions can be found in Supplementary Table 1. Following each PCR, amplicons were purified using AMPure XP beads (Beckman Coulter) at a 0.7:1 ratio. Libraries were quantified using Qubit (Invitrogen) and analyzed using an Agilent 4200 TapeStation. Combined libraries were loaded at 5pM with 25% PhiX control (Illumina) and sequencing was performed with a v2 Nano Reagent kit 2×250bp on a MiSeq machine at the Genomic Research Unit (GRU), Tel Aviv University. For Off-target analysis, raw fastq files were submitted to Fast Length Adjustment of Short Reads (FLASH) (https://github.com/ebiggers/flash)^28^. The default parameters were changed to allow for lower max mismatch density ratio of 0.1. The resultant files were submitted to CRISPRpic (https://github.com/compbio/CRISPRpic)^29^, with a wider mutagenic window of 10 bp on either side of the DNA double-strand breaks. Presented data pools all mutation types detected.

For SHM and selection analysis, r aw fastq files were submitted to FLASH using the default parameters. The resultant files submitted to Bowtie2 alignment analysis (https://github.com/BenLangmead/bowtie2)^30^ compared to the engineered 3BNC117 sequence, using local mode and the “xeq” parameter for match and mismatch annotations. Using a specific script, unaligned reads were filtered, as well as reads not within 80-115% of the original length and reads with more than 15% mutated bases. The primer annealing sites at both ends of the sequences were omitted from the analysis. All bases considered as mutated in this analysis had a Q score higher than 20. The Selecton software^31^ was used to run M8 and M8a models in order to infer positive selection and likelihood ratio test was performed between the null model (M8a) and the alternative model (M8) to determine which model better fits the data. All P-values were corrected for multiple testing using false discovery rate (FDR)^32^. All alignments and phylogenies supported the M8 alternative model where positive selection is enabled.

### CHANGE-seq

Genomic DNA from spleens of wild type C57BL/6OlaHsd using Gentra PureGene Tissue Kit (Qiagen) and quantified using Qubit (Invitrogen) according to manufacturer instructions. CHANGE-seq was performed as previously described^20^. Briefly, purified genomic DNA was tagmented with a custom Tn5-transposome to an average length of 400 bp, followed by gap repair with Kapa HiFi HotStart Uracil+ DNA Polymerase (KAPA Biosystems) and Taq DNA ligase (NEB). Gap-repaired tagmented DNA was treated with USER enzyme (NEB) and T4 polynucleotide kinase (NEB). Intramolecular circularization of the DNA was performed with T4 DNA ligase (NEB) and residual linear DNA was degraded by a cocktail of exonucleases containing Plasmid-Safe ATP-dependent DNase (Lucigen), Lambda exonuclease (NEB) and Exonuclease I (NEB). *In vitro* cleavage reactions were performed with 125 ng of exonuclease-treated circularized DNA, 90 nM of EnGen® Sau Cas9 protein (NEB), NEB buffer 3.1 (NEB) and 270 nM of sgRNA (Synthego), in a 50 μl volume. Cleaved products were A-tailed, ligated with a hairpin adaptor (NEB), treated with USER enzyme (NEB) and amplified by PCR with barcoded universal primers NEBNext Multiplex Oligos for Illumina (NEB), using Kapa HiFi Polymerase (KAPA Biosystems). Libraries were quantified by qPCR (KAPA Biosystems) and sequenced with 151 bp paired-end reads on an Illumina MiniSeq instrument. CHANGE-seq data analyses were performed using open-source CHANGE-seq analysis software (https://github.com/tsailabSJ/changeseq).

### ELISA

High binding microplates (Greiner Bio-One) were coated with 2 μg/ml of an anti-idiotipic antibody against 3BNC117 in PBS overnight at 4°C. Plates were washed with PBST, blocked for an hour with 5% BSA in PBST and washed again. For 3BNC117 IgG quantification, samples were diluted 1:50-500 fold and a standard was made using purified 3BNC117 serially diluted in PBS. For 3BNC117 isotype detection, samples were serially diluted as described in the figures. Samples and standards were incubated for an hour. Plates were then applied with HRP conjugated detection antibodies: anti-mouse IgA (Abcam), anti-mouse IgG, anti-mouse IgG1, anti-mouse IgM (Jackson ImmunoResearch) or anti mouse IgG2c (Bio-Rad Laboratories) at 2 μg/ml in PBST and were incubated for another hour. A list of antibodies used in these experiments may be found in Supplementary Table 1. Before detection with QuantaBlu (ThermoFisher) according to manufacturer protocol, plates were washed for an additional round. Detection was done in a Synergy M1 Plate reader (Biotek). When absolute quantitation is presented, the concentration of 3BNC117 was determined by reference to the dilution factor of the standard curve.

### Neutralization Assays

Under sterile BSL2/3 conditions, the PSG3 plasmid was co-transfected into HEK293T cells along with JRFL or YU2 HIV envelope plasmids using Lipofectamine 2000 transfection reagent (ThermoFisher Scientific) to produce single-round of infection competent pseudo-viruses representing multiple clades of HIV. 293T cells were plated in advance overnight with DMEM medium + 10% FBS + 1% Pen/Strep + 1% L-glutamine. Transfection was done with Opti-MEM transfection medium (Gibco) using Lipofectamine 2000. Fresh medium was added 12 h after transfection. Supernatants containing the viruses were harvested 72 h later. In sterile 96-well plates, 25 μl of virus was immediately mixed with 25 μl of serially diluted (2×) protein A/G purified IgG (ThermoFisher) from mouse sera (starting at 500 μg/ml) and incubated for one hour at 37 °C to allow for antibody neutralization of the pseudoviruses. 10,000 TZM-bl cells/well (in 50 μl of media containing 20 μg/ml Dextran) were directly added to the antibody virus mixture. Plates were incubated at 37 °C for 48 h. Following the infection, TZM-bl cells were lysed using 1× luciferase lysis buffer (25 mM Gly-Gly pH 7.8, 15 mM MgSO4, 4 mM EGTA, 1% Triton X-100). Neutralizing ability disproportionate with luciferase intensity was then read on a Biotek Synergy 2 (Biotek) with luciferase substrate according to the manufacturer’s instructions (Promega).

### qPCR

For copy number quantification of the donor AAV in tissue samples, DNA was extracted using DNeasy Blood & Tissue Kit (Qiagen) with RNAse treatment on-column. Each sample was analyzed for both internal control (Albumin intron^26^) and the donor AAV. For quantification of donor AAV copy number per haploid genome, a standard curve was used. Standards were prepared from a PCR PrimeStar MAX (Takara) reaction using naive C57BL/6OlaHsd mice genomic DNA for the internal control or donor AAV plasmid for the Donor sample and purified using AMPure XP beads (Beckman Coulter) at a 1:1 ratio. A list of primers used for Donor and internal control reactions can be found in Supplementary Table 1. Standard curve amplicons were quantified using Qubit (Invitrogen) and serially diluted 8 times. For AAV Cas9 quantification RNA was extracted from tissues using RNeasy Mini Kit (Qiagen) with DNAse treatment on-column and post-purification using RQ1 DNAse (Promega). Reverse transcription was performed using RevertAid (ThermoFisher) and random hexamer primers. Data collection and analysis were performed on a StepOnePlus qPCR System (Applied Biosystems) using SYBRGreen (PCR Biosystems). For fold change of AAV titers and Cas9 relative expression, we used the relative quantity method^33^.

### Flow Cytometry

Harvested cells from spleen, bone marrow or blood were resuspended in cell staining buffer (Biolegend) and incubated with 2μg/100μl of human anti-3BNC117 for 10 mins, washed and resuspended again in cell staining buffer containing primary antibodies. Secondary staining was performed in the dark, for 15 mins, with anti-human IgG1 AF647 (Abcam) or anti-human IgK BV421 (Biolegend). A list of antibodies and respective dilutions used in these experiments can be found in Supplementary Table 1. Then, cells were washed and data acquisition was performed on a CytoFLEX (Beckman Coulter) or FACS Aria III (BD Biosciences) for experiments involving cell sorting. Data were compiled and analyzed using Kaluza Analysis 2.1 (Beckman Coulter). Gating strategy can be found in Fig. S19.

### Nucleic Acid Manipulations

For Reverse Transcription PCR demonstrating 3BNC117 gene integration into the IgH locus, RNA was extracted from sorted engineered B cells (3BNC117^+^, CD4^-^, CD19^+^) on a FACS BD AriaIII (BD Biosciences) using the RNeasy Mini Kit (Qiagen) with DNAse treatment on-column and reverse transcription was performed using RevertAid (ThermoFisher) and Oligo dT primers. PCR on the resulting cDNA was performed for 35 cycles using PrimeStar MAX (Takara). Then, a semi-nested PCR was performed using PrimeStar MAX (Takara) for 35 cycles. A list of primers used for these reactions may be found in Supplementary Table 1. Following each PCR, resulting amplicons were analyzed by Agarose gel electrophoresis as compared to a standardizing ladder (Hylabs, GeneDireX 1Kb plus DNA ladder RTU or 100bp DNA ladder H3 RTU) and total reactions were purified using AMPure XP beads (Beckman Coulter) at a 1:1 ratio. Purified amplicons were Sanger sequenced at the DNA Sequencing Unit, Tel Aviv University and presented alignment of the chromatograms was performed using SnapGene (GSL Biotech).

For TIDE analysis of on target cleavage, genomic DNA from tissues was extracted using Gentra PureGene Tissue Kit (Qiagen). PCR was performed for 35 cycles using PrimeStar MAX (Takara). Primers used for these reactions can be found in Supplementary Table 1. For the control samples, three independent PCR were performed on independent genomic DNA samples, collected from splenic tissue of naive C57BL/6OlaHsd mice. Resulting amplicons were purified using AMPure XP beads (Beckman Coulter) at a 1:1 ratio. Purified amplicons were Sanger sequenced at the DNA Sequencing Unit, Tel Aviv University. For each sample, multiple sequencing reactions were performed using either primers. Then, samples were compared using TIDE (https://tide.nki.nl/)^34^. For control samples, we performed reciprocal sample comparisons from the independent initial PCR reactions.

### Statistics

Statistical analyses were performed on distinct samples using Prism (GraphPad). For Area under the curve, in each group, the mean AUC and SD was calculated and these values were compared by t-test. All t-tests were performed as two tailed. For the TIDE analysis, each comparison between a control sample and an independently produced PCR reaction from 3 independent mice were used. Each Figure legend denotes the statistic used, central tendency and error bars.

## DATA AVAILABILITY

Data is available in the main text, in the Supplementary Data and Materials. Illumina sequencing data can be accessed in the SRA database under accession code PRJNA706552. The authors declare that unique materials used are readily available from the authors upon MTA agreement.

## SUPPLEMENTARY FIGURES

**Fig. S1:**
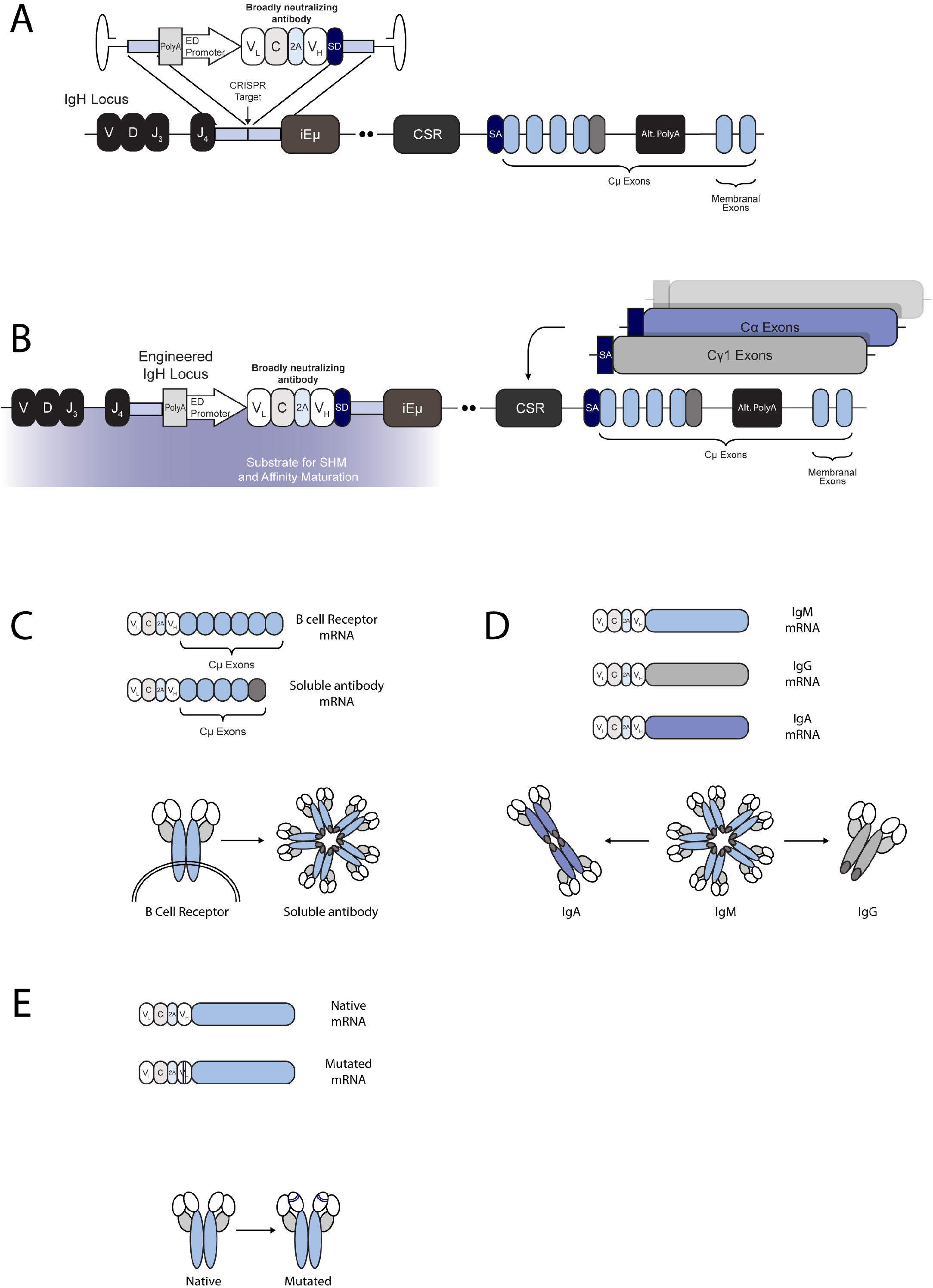
Targeting an antibody to the IgH locus of B cells allows for antigen induced activation, SHM, CSR and affinity maturation. **A**. Engineering scheme depicting the integration site of the 3BNC117 cassette into the J-C intron of the IgH locus. 3BNC117 is targeted using CRISPR/Cas9 downstream of the last J segment (J_4_) and upstream of the intronic enhancer (iEμ), class switch recombination locus (CSR) and the IgH Cμ exons. **B**. The integrated antibody may undergo somatic hyper mutation (SHM) and CSR. Indicated below is the area targeted by AID for SHM. The arrow indicates a CSR event where the IgHCμ exons will be replaced by exons coding for another constant domain. For the IgHCμ exons we indicate the alternative polyA, active when the antibody is secreted, and the membranal exons, expressed when the polyA is not active. **C**. The bNAb mRNA is terminated by alternative poly adenylation sites allowing for membranal (BCR) or soluble expression, before and after differentiation into a plasma cell, respectively. mRNA (above) and protein (below) scheme of a BCR or soluble antibody. **D**. The donor AAV does not code for the constant domain, rather it uses a splice donor (SD) to splice with the endogenous constant. Therefore, when an engineered cell undergoes CSR the integrated antibody may be expressed as a different isotype. mRNA (above) and protein (below) scheme of the antibody with different classes. **E**. The bNAb is targeted to an AID active locus. Therefore, SHM in the antibody coding genes may allow for clonal expansion due to affinity maturation. mRNA (above) and protein (below) scheme of the antibody undergoing SHM.

**Fig. S2:**
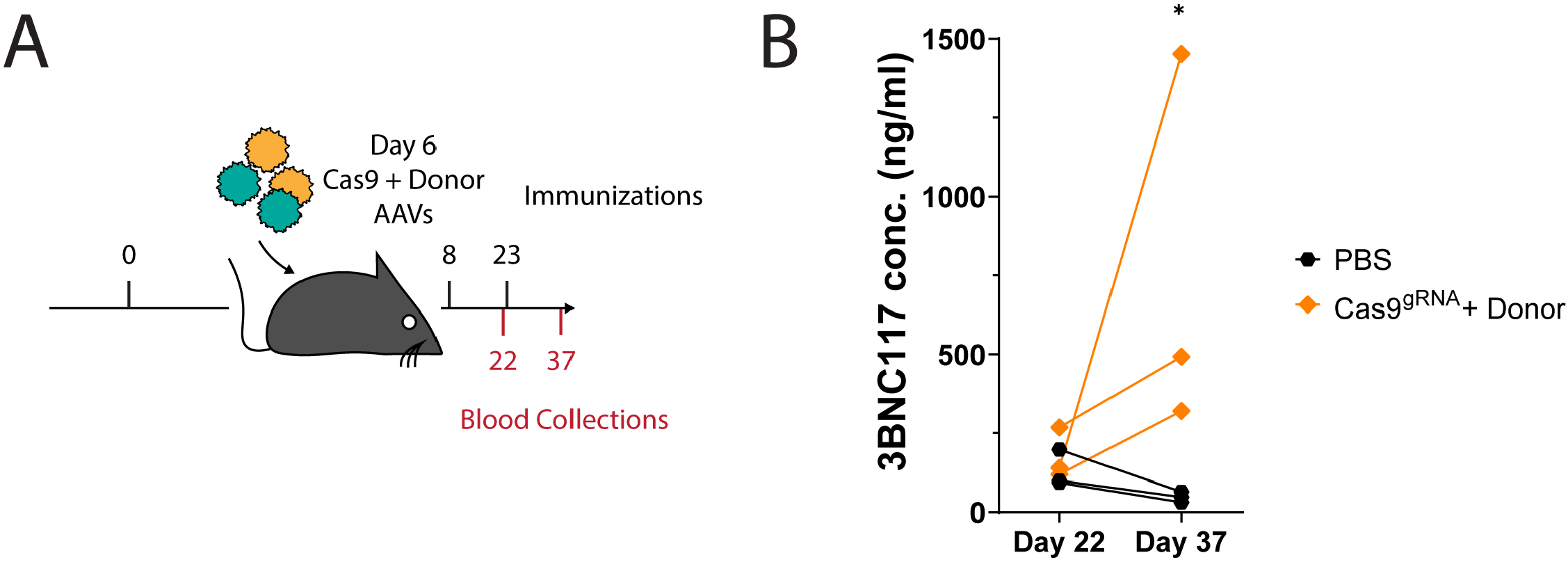
*In vivo* engineered B cells are activated by the BG505 MD39 HIV antigen. 3BNC117 IgG titers as quantified by ELISA, using an anti-idiotypic antibody as compared to a PBS injected group, similarly immunized with gp120. * = pv<0.05 matched, two-way ANOVA with Šidák’s multiple comparison. In this figure the PBS group is is the same as described in Fig. 4C.

**Fig. S3:**
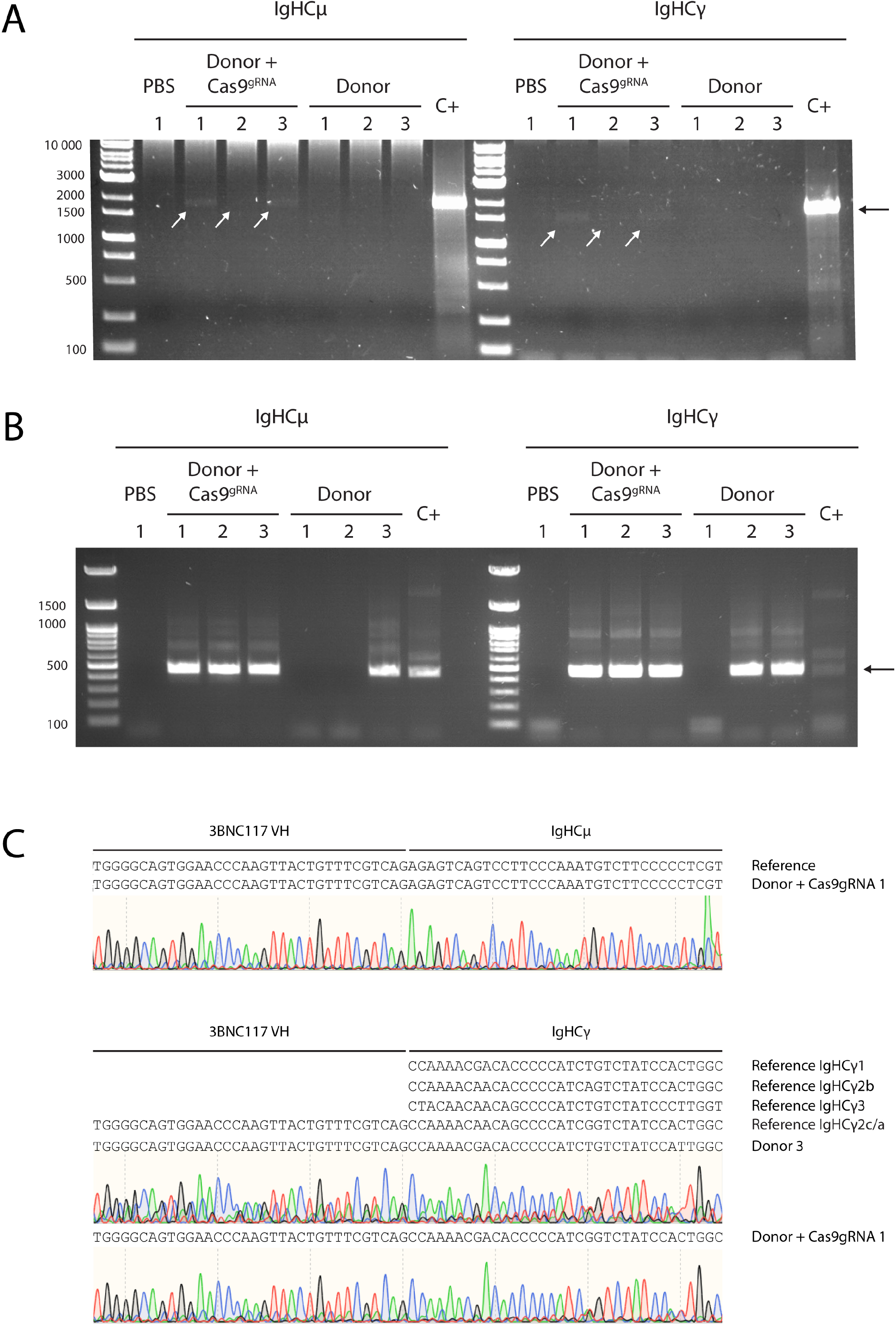
Validation of on-target integration. **A**. RT-PCR on RNA from sorted, 3BNC117^+^, CD19^+^, CD4^-^ blood lymphocytes from day 37. Here, we used a reverse primer in a membranal exon of either IgHCμ or IgHCγ (all subtypes) and a forward primer on the V_H_ of the coded 3BNC117. Numbers indicate different mice, injected with either a) PBS, b) The donor vector and the CMV-Cas9^gRNA^ vector, or c) the donor vector only, as indicated above the gels. Control sample comes from *in-vitro* engineered primary mouse splenic lymphocytes. Ladder sizes are indicated on the left. Arrow indicates the expected amplicon size. **B**. Total DNA from the previous reaction was purified and a semi-nested PCR with the same forward primer and a reverse primer on the CH1 of the constant domains. Ladder sizes are indicated on the left. Arrow indicates the expected amplicon size. **C**. Sanger sequencing alignment and chromatogram of the purified amplicon from the previous step. Reference sequences are indicated above. For the IgHCγ, each subtype reference is indicated.

**Fig. S4:**
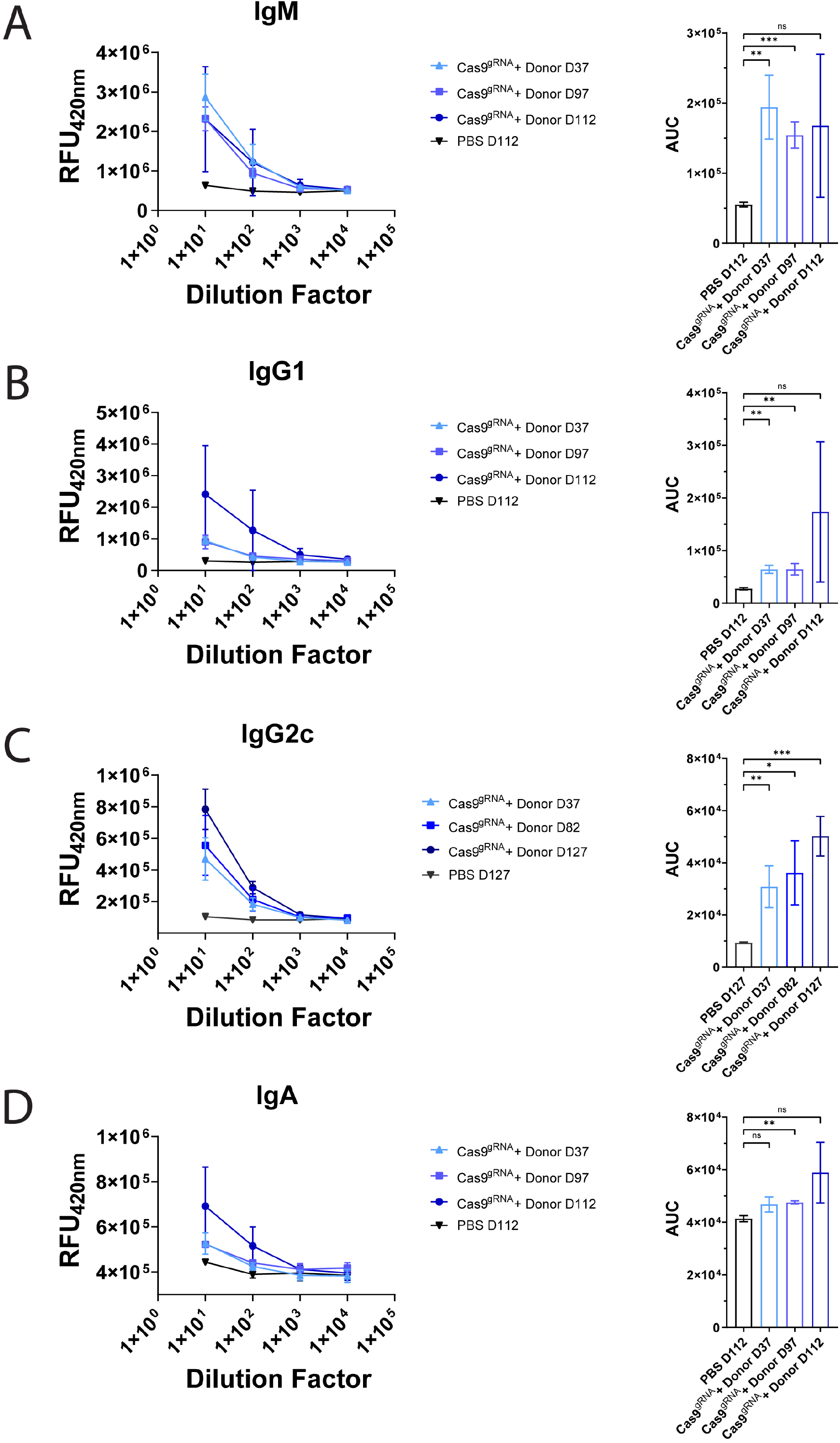
Multiple isotypes of the 3BNC117 antibody are expressed by engineered B cells. **A-D**. ELISA for each isotype, with the AUC comparison on the right. A. IgM, B. IgG1, C. IgG2c and D. IgA. All samples come from the CMV-Cas9^gRNA^ + Donor injected mice, at different time points as indicated in each legend. Mean and SD are indicated. For AUC, * = pv<0.05, ** = pv<0.01, *** = pv<0.001, unpaired t-test. n=3.

**Fig. S5:**
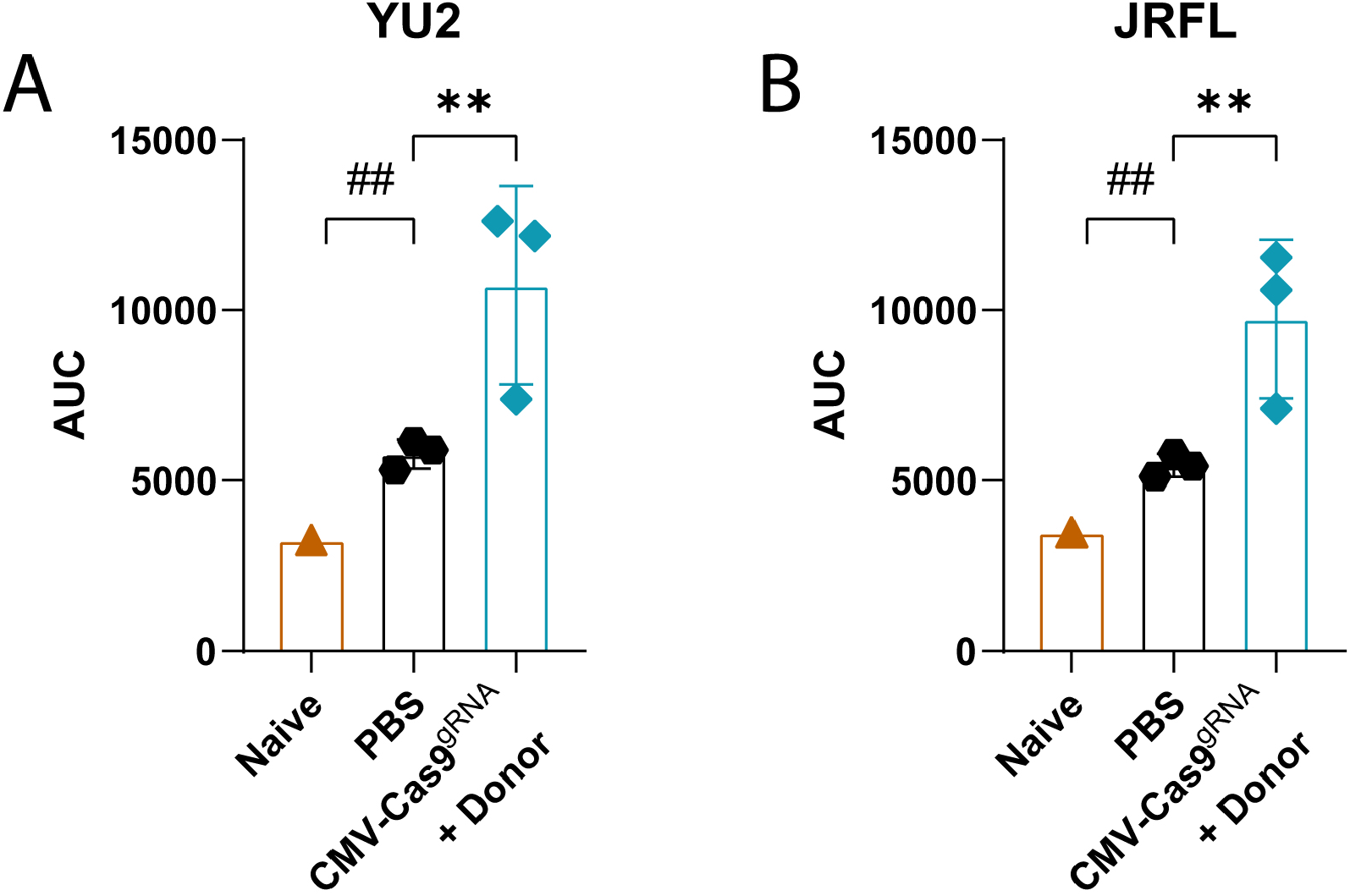
Area under the Curve (AUC) of Fig. 1E for YU2.DG **A**. and JRFL **B**. ** = pv<0.01with unpaired t-test for CMV-Cas9gRNA + Donor to PBS comparison and ## = pv<0.01 with one-sample t-test for Naïve to PBS comparison. n=3 for CMV-Cas9gRNA + Donor and PBS. Naïve sample from a single, non-immunized, non-AAV-injected mouse.

**Fig. S6:**
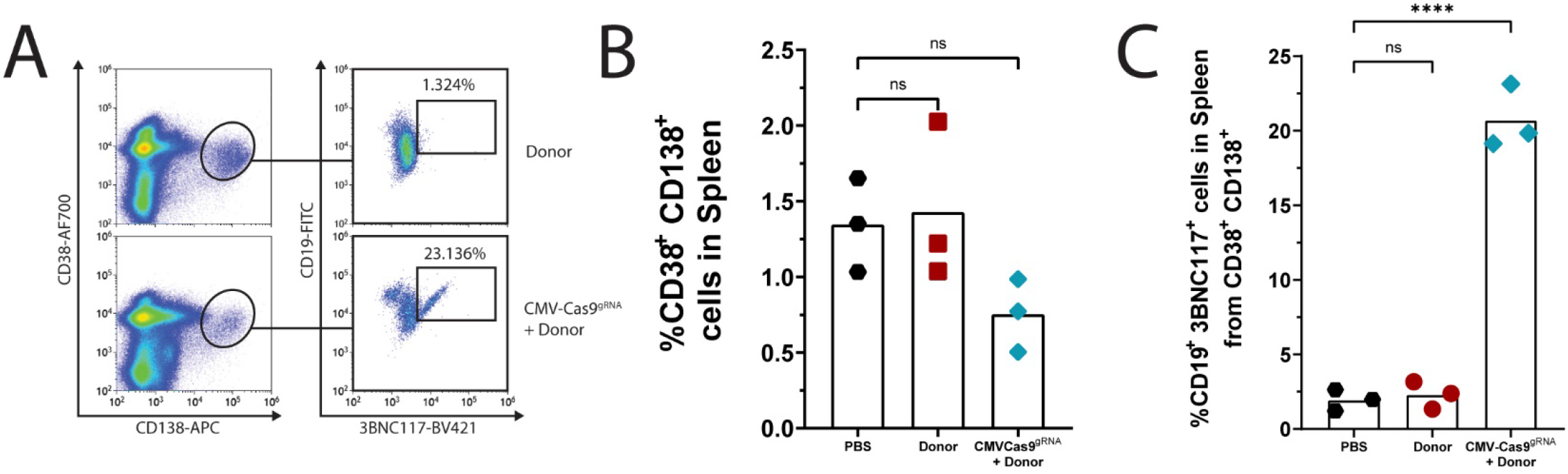
A. Flow cytometry plots demonstrating 3BNC117 expression among plasmablasts (CD38^+^, CD138^+^, CD19^+^) in the spleen at day 136. **B**. Quantification of A for total plasmablasts (CD38^+^ CD138^+^). **C**. Quantification of A. for engineered plasmablasts (CD38^+^ CD138^+^ 3BNC117). Mean is indicated by the bars. ns= non-significant, **** = pv < 0.0001, One-way ANOVA with Tukey’s multiple comparison. Pregating for A. is live, singlets.

**Fig. S7:**
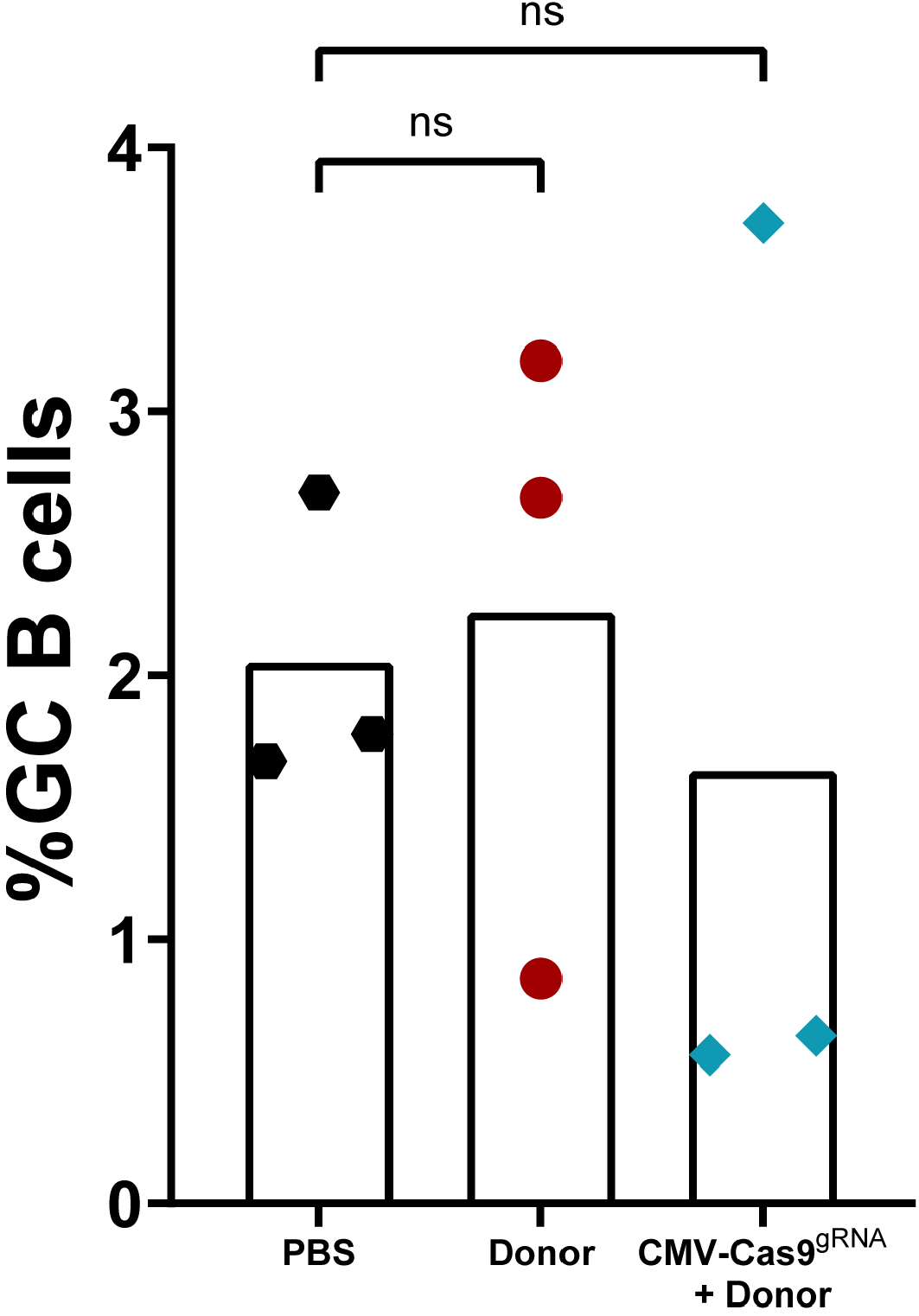
Quantification by flow cytometry of total GL7^+^ Fas^+^ GC B cells in spleens at day 136. Mean is indicated by the bars. ns= non-significant, One-way ANOVA with Tukey’s multiple comparison.

**Fig. S8:**
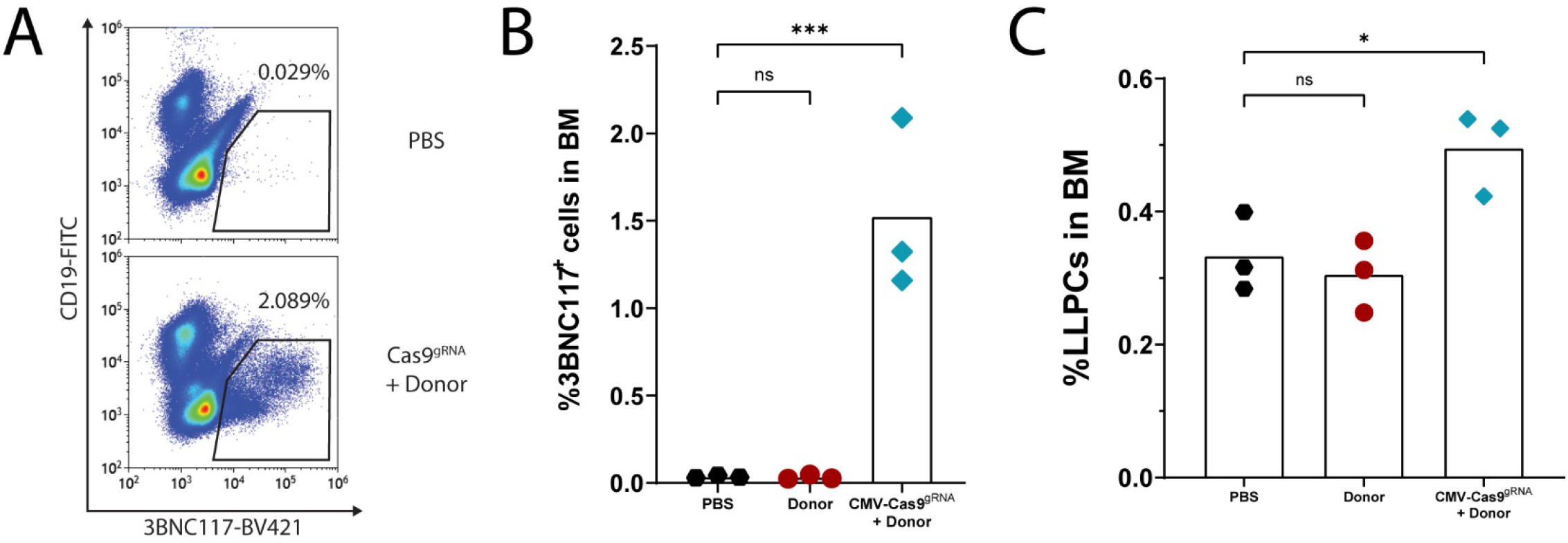
**A**. Flow cytometry plots demonstrating the presence of 3BNC117-expressing B cells in the bone marrow. **B**. Quantification of A. **C**. Quantification of total long-lived plasma cells (LLPCs) (CD19^low^ CD138^+^). Mean is indicated by the bars. ns= non-significant, * = pv < 0.05, *** = pv < 0.001 One-way ANOVA with Tukey’s multiple comparison.

**Fig. S9:**
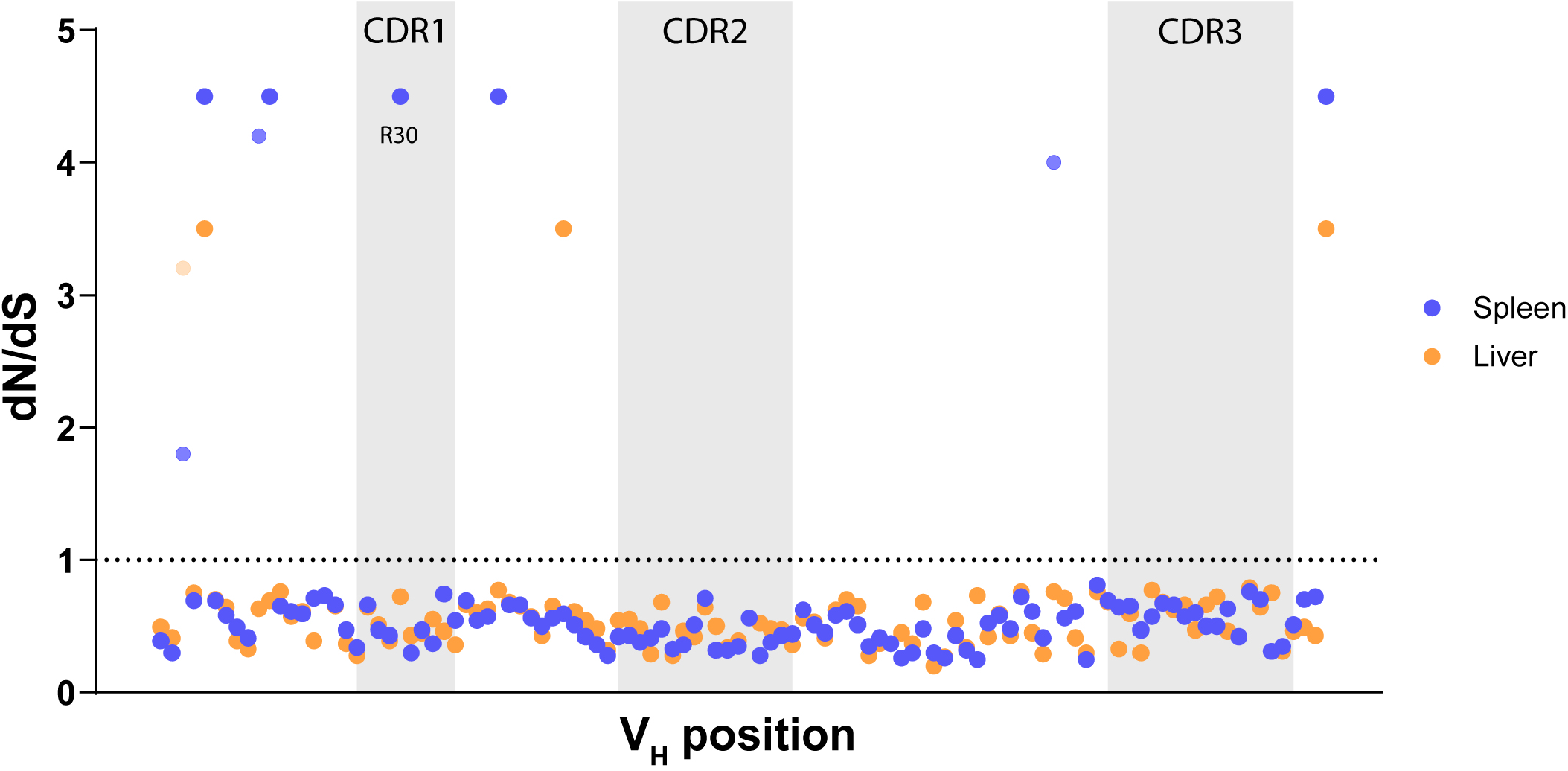
dN/dS values for the positions along the V_H_ segment based on Illumina sequencing of DNA amplified from the spleen (blue) or liver (orange) of a single mouse. Light shading indicates lower boundary value <1. Grey shading indicates CDR loops. The R30 position is indicated.

**Fig. S10:**
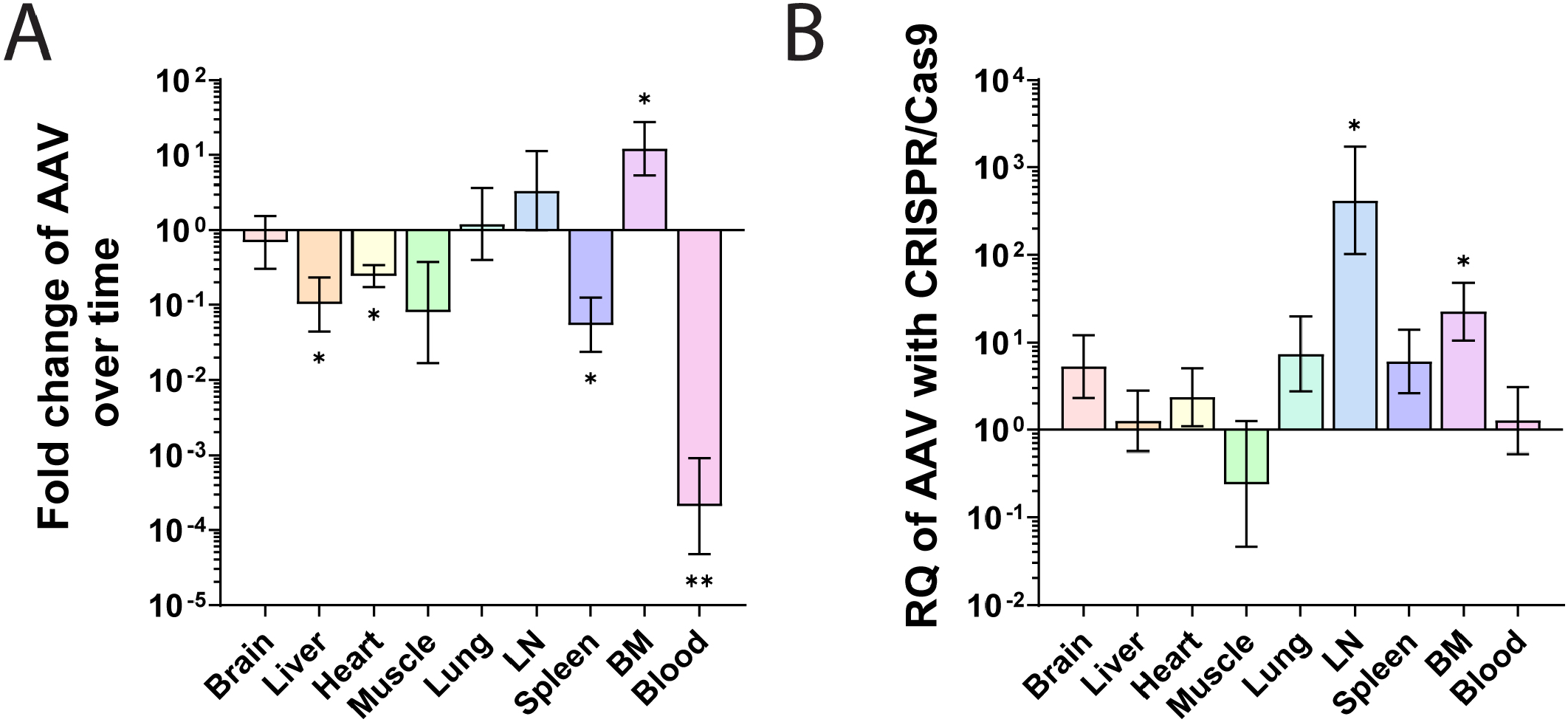
**A**. Fold change of donor AAV copy number over time, as determined by qPCR. Mean with upper and lower boundaries are indicated. * = pv<0.05 and ** = pv<0.01 for unpaired t-test comparing the two time points, day 37 and day 136. **B**. Relative quantity of Donor AAV in the CMV-Cas9^gRNA^ + donor group as compared to the donor only group, at day 136. Mean with upper and lower boundaries are indicated. n=3. For both A. and B. data presented includes samples presented in Fig. 3B-C and samples from additional tissues.

**Fig. S11:**
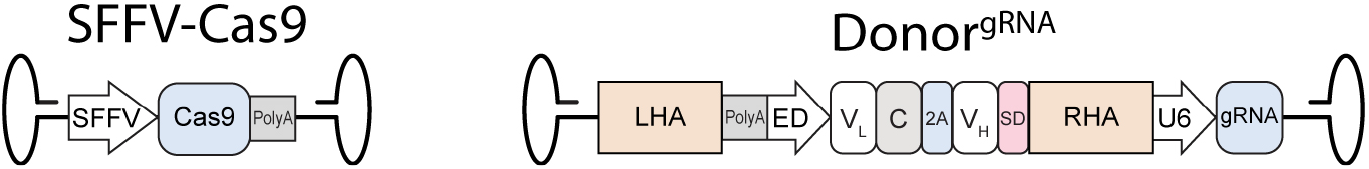
Vector maps of the AAVs coding for the Donor^gRNA^ and the SFFV-Cas9.

**Fig. S12:**
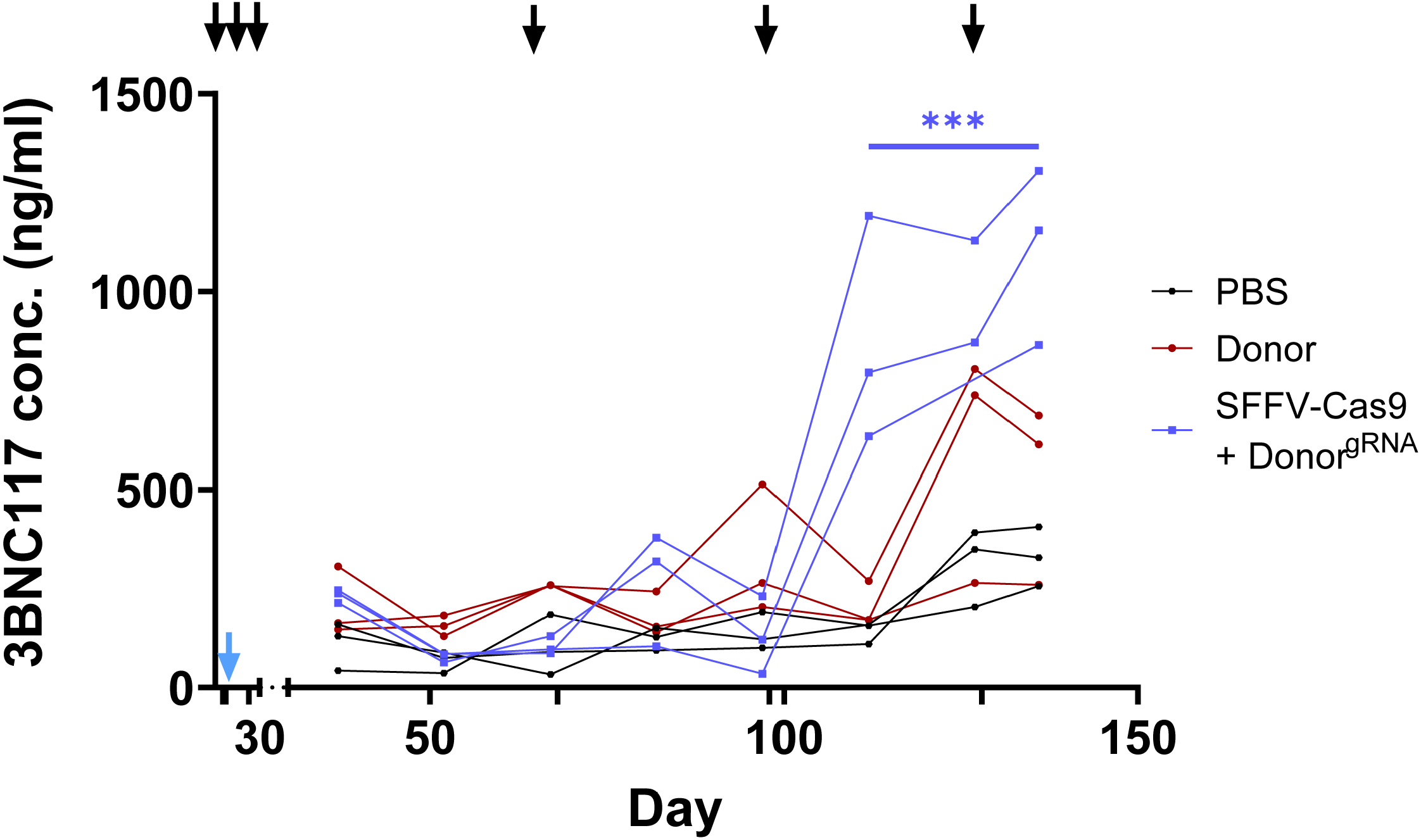
3BNC117 IgG titers as quantified by ELISA over time in the SFFV-Cas9 + Donor^gRNA^ group. The black arrows indicate immunizations and the blue arrow indicates AAV injection. Each mouse is indicated. *** = pv<0.001 for two-way ANOVA comparing the SFFV-Cas9 + Donor^gRNA^ group to the Donor group. n=3. In this figure, the PBS and Donor control groups are the same as for Fig. 1C.

**Fig. S13:**
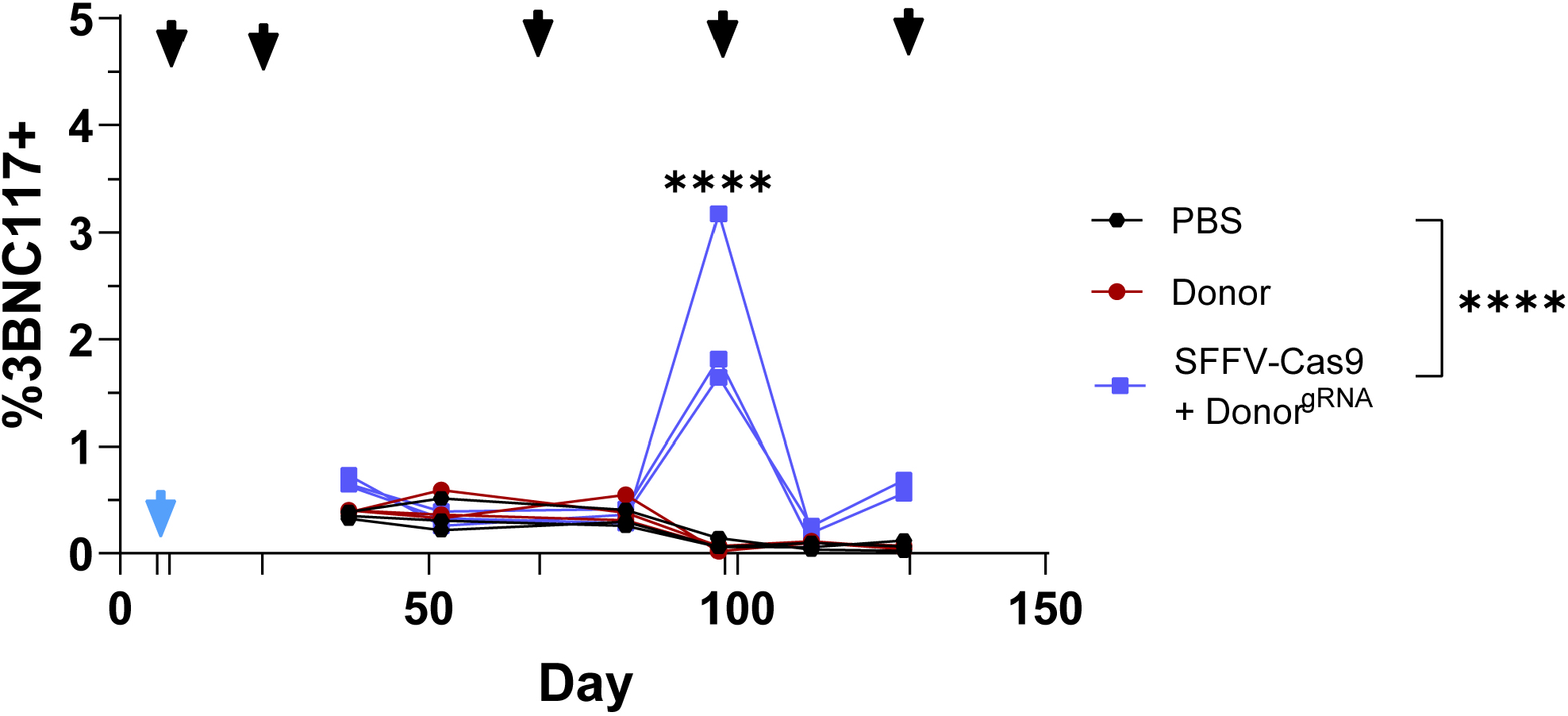
3BNC117^+^, CD19^+^, CD4^-^ blood lymphocytes as followed by flow cytometry over time in the SFFV-Cas9 + Donor^gRNA^ group. The black arrows indicate immunizations and the blue arrow indicates AAV injection. **** = pv<0.0001 Two-way ANOVA with Šidák’s multiple comparison for time points comparison to PBS. n=3. In this figure, the PBS and Donor control groups are the same as for Fig. 2B.

**Fig. S14:**
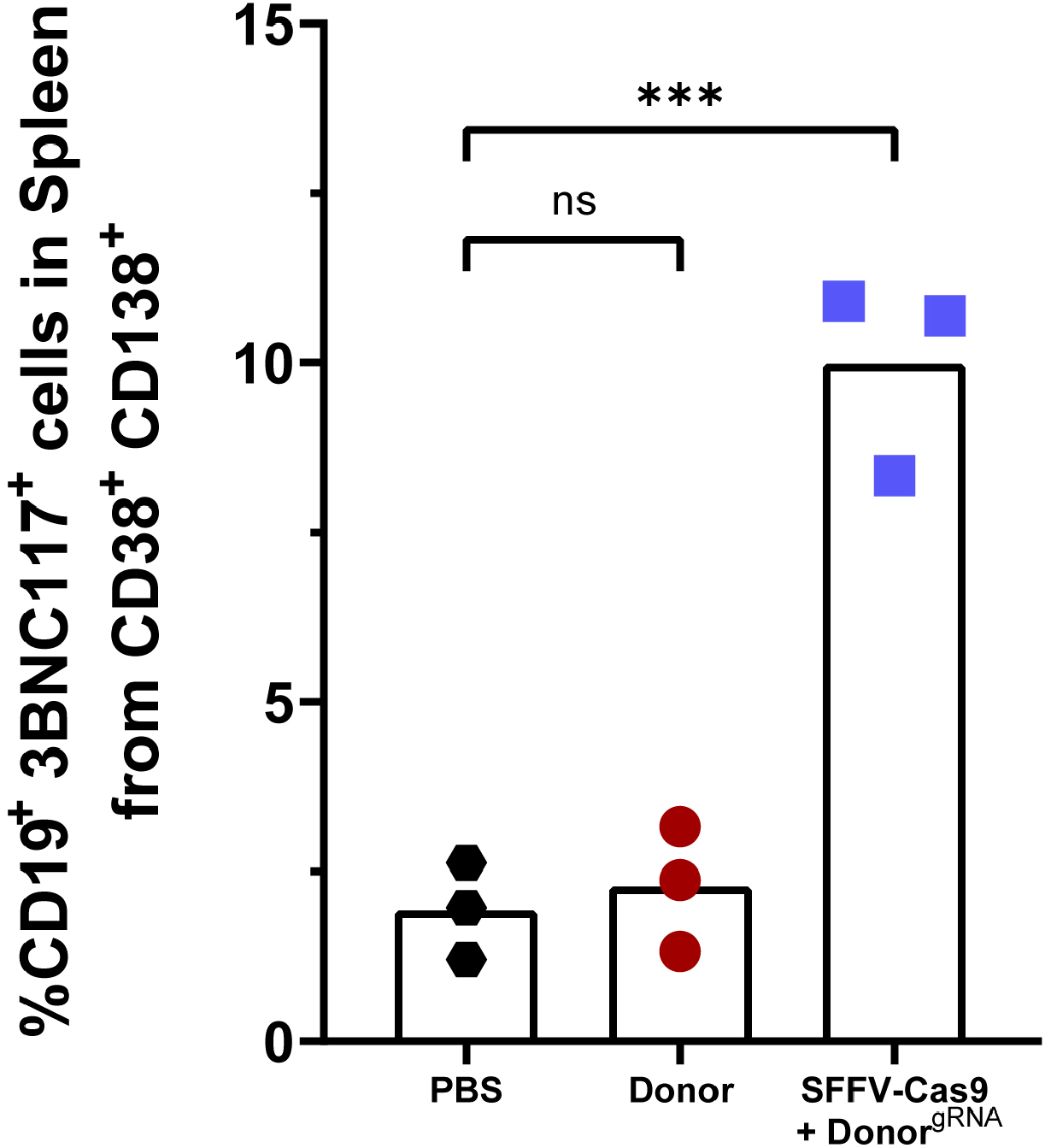
Quantification by flow cytometry of 3BNC117 ^+^, CD19^+^, CD38^+^, CD138^+^ plasmablasts in the spleens of the SFFV-Cas9 + donor^gRNA^ group at day 136. Mean is indicated by the bars. *** = pv<0.001, one-way ANOVA with Tukey’s multiple comparison. In this figure, the PBS and Donor control groups are the same as for Fig. S6.

**Fig. S15:**
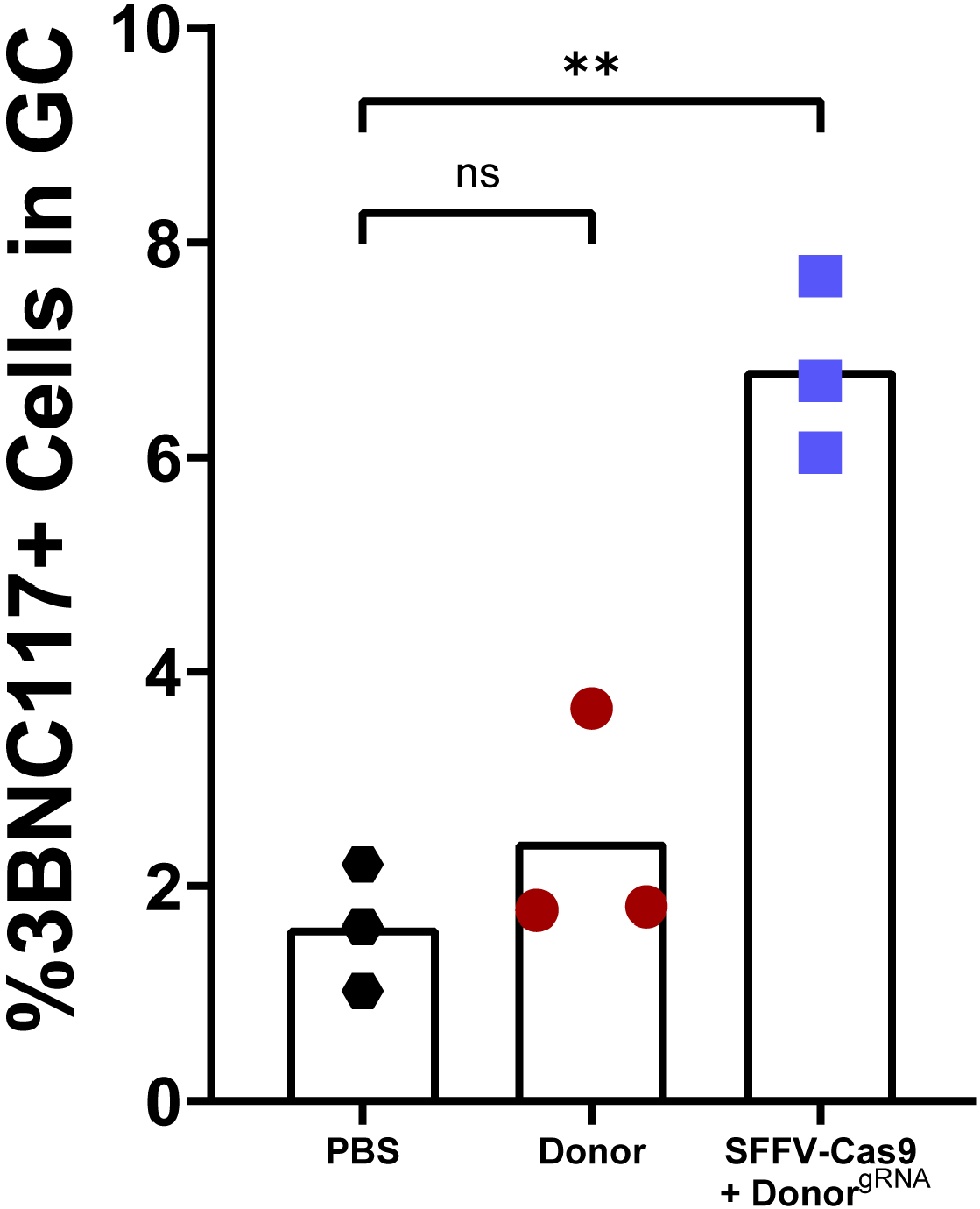
Quantification by flow cytometry of GL7^+^, Fas/CD95^+^ GC B cells in the spleens of the SFFV-Cas9 + Donor^gRNA^ group at day 136. Mean is indicated by the bars. ** = pv<0.01, one-way ANOVA with Tukey’s multiple comparison. In this figure, the PBS and Donor control groups are the same as for Fig. 2E.

**Fig. S16:**
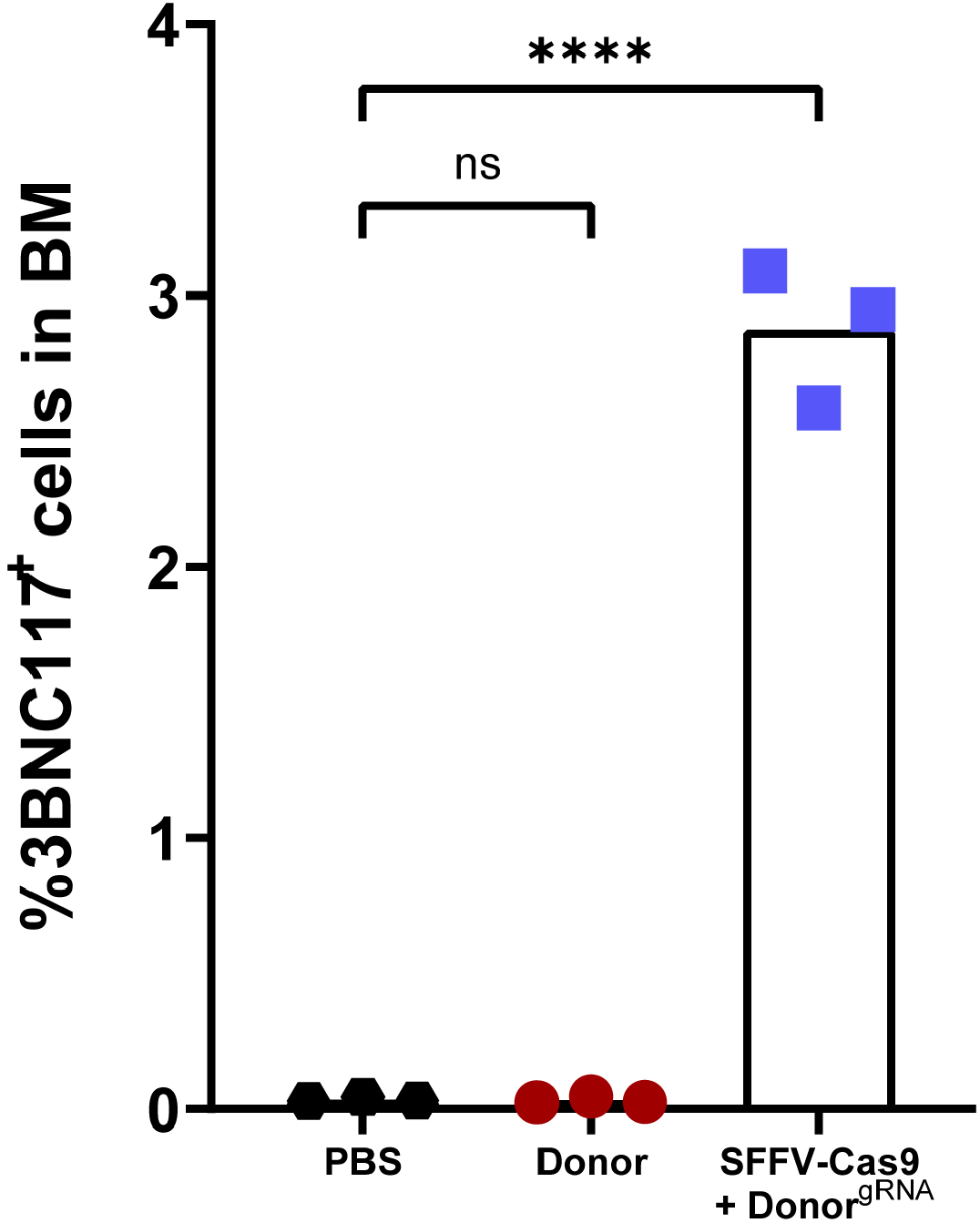
Quantification by flow cytometry of 3BNC117^+^ cells in total bone marrow (BM) of the SFFV-Cas9 + Donor^gRNA^ group at day 136. Mean is indicated by the bars. **** = pv<0.0001, one-way ANOVA with Tukey’s multiple comparison. In this figure, the PBS and Donor control groups are the same as for Fig. S8B.

**Fig. S17:**
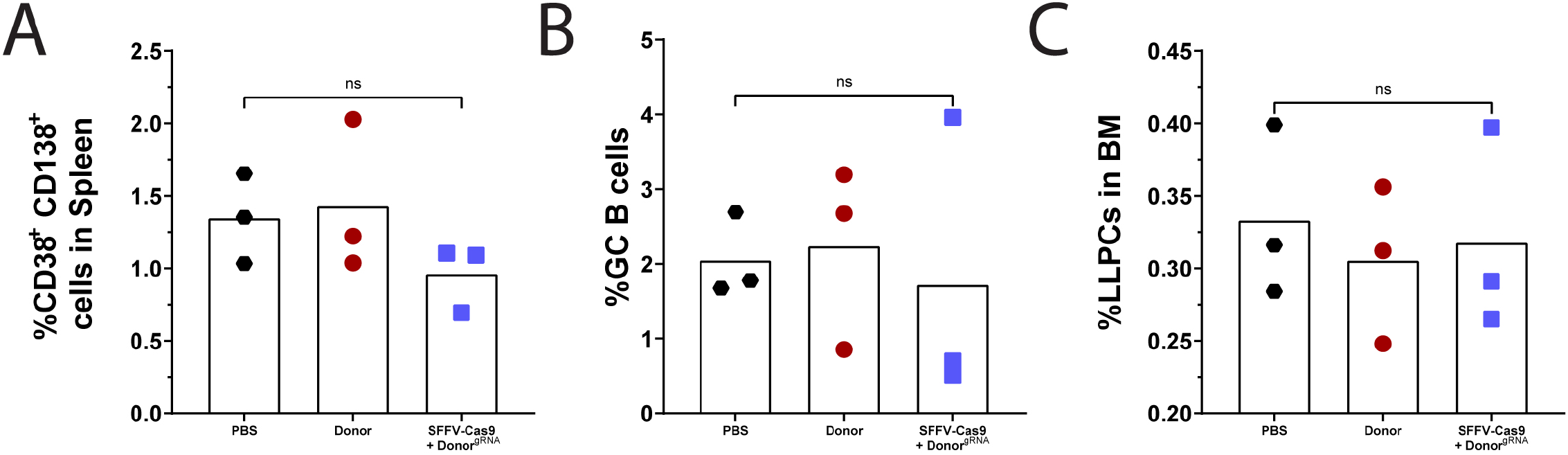
Assessing overall immune homeostasis. Quantification by flow cytometry of total CD38^+^ CD138^+^ plasmablasts in spleen (**A.)**, total GL7^+^, Fas^+^ GC B cells in the spleen (**B.)** and total CD19^low^, CD138^+^ long-lived plasma cells (LLPCs) in bone marrow (**C.)**, at day 136. Mean is indicated by the bars. ns = non-significant, one-way ANOVA with Tukey’s multiple comparison. In this figure, the PBS and Donor control groups are the same as for Fig. S6B, S7, S8C.

**Fig. S18:**
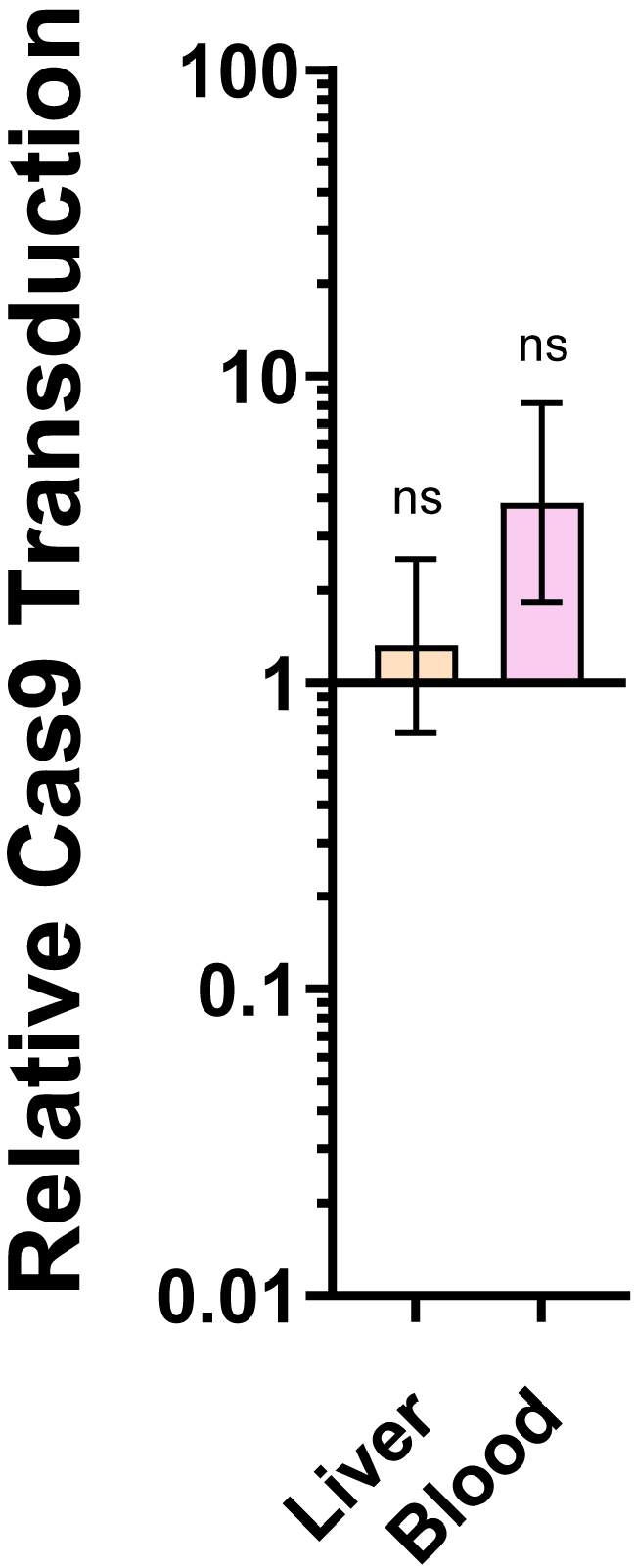
Relative copy number of the saCas9 coding AAV, entailing either the CD19 or SFFV promoters, in liver and blood. Error bars correspond to lower and upper boundaries derived from unpaired t test for A. and B., ns = non-significant. n=3.

**Fig. S19:**
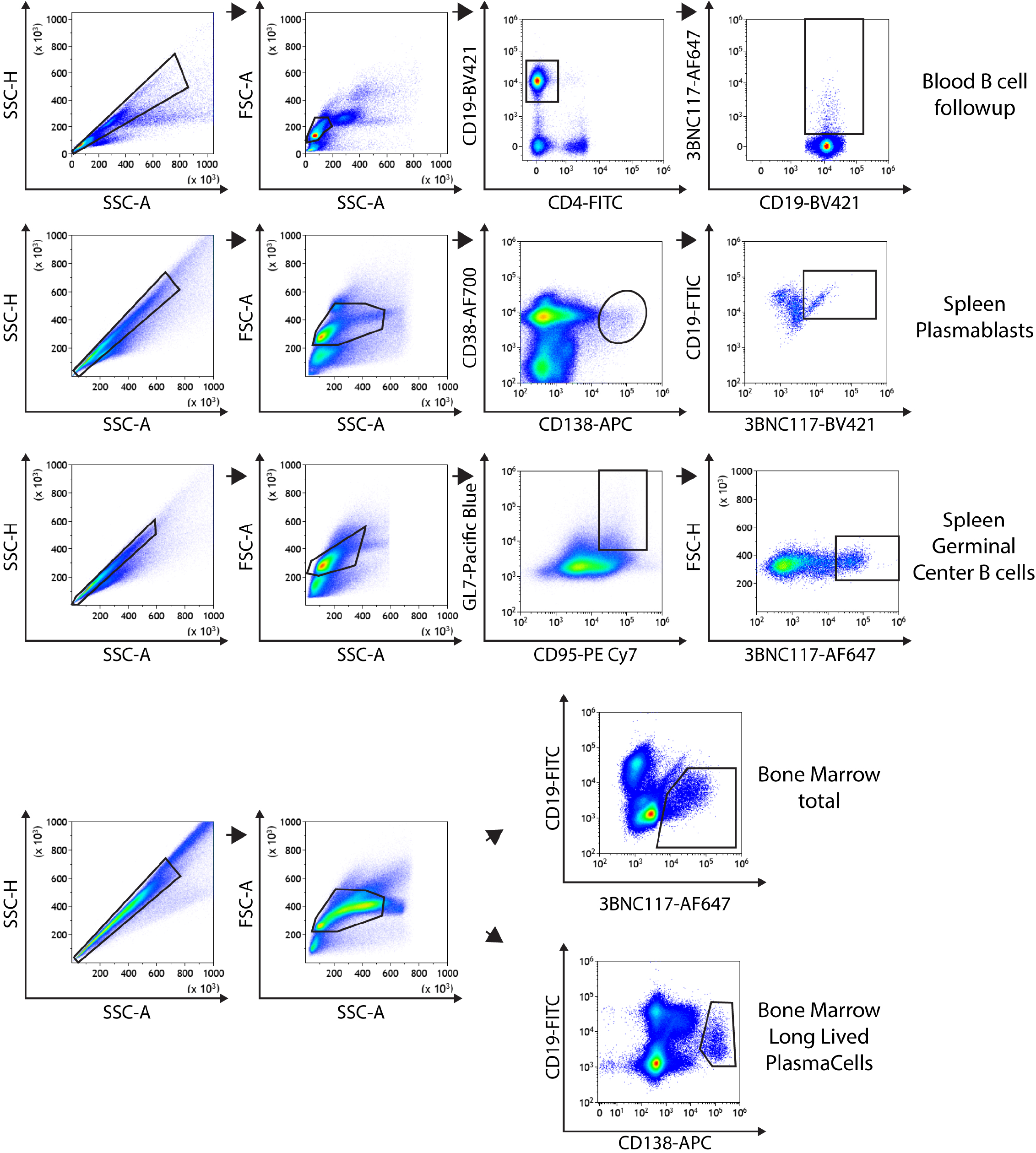
Gating strategy for each experiment in this study.

## ACKNOWLEDGEMENTS

We thank the Veterinary Service Center, Tel Aviv University for animal husbandry. The IDRFU, GRU and SICF units, Tel Aviv University for logistic support and council. We also thank Leeor Vardi, Maoz Gelbart, Hila Kobo, David Burstein, Itai Benhar, Natalia Freund, Mark Kay, Tal Akriv, Daniel Nataf and Natalia Gritsenko for reagents and feedback. This research was funded by the H2020 European Research Council grant 759296 570 (A.B.) and the Israel Science Foundation grants: 1632/16 (A.B.), 2157/16 (A.B.), The Bill and Melinda Gates Foundation: OPP1183956 (J.E.V.), National Institutes of Health: R01 AI128836 and R01 AI073148 (D.N.), Edmon J. Safra Center for Bioinformatics fellowship (T.K. and A.S.), St. Jude Children’s Research Hospital and ALSAC, National Institutes of Health (NIH) Office Of The Director (OD) Somatic Cell Genome Editing (SCGE) initiative grant U01AI157189 (S.Q.T.). The content is solely the responsibility of the authors and does not necessarily represent the official views of the National Institutes of Health.

## AUTHOR CONTRIBUTIONS

A.D.N designed, performed and analyzed the study; C.R.L. performed CHANGE-seq; S.Q.T. supervised CHANGE-seq experiments; N.Z. and T.K. performed bioinformatical analyses; A.S. and R.R.A supervised the bioinformatical analyses; N.Z., M.H.F. and I.R. helped with sample processing; M.T. and D.H. performed neutralization assays; D.N. and J.E.V. supervised neutralization assays; I.D. contributed to supervising the study; A.D.N. and A.B. drafted and revised the manuscript; A.B. Conceptualized and supervised the study.

